# Brain pericytes exhibit spatially organized and dynamically regulated molecular heterogeneity

**DOI:** 10.64898/2026.08.17.745290

**Authors:** Sheng-Fu Huang, Lukas Glandorf, Sidonie Sauvageot, Chaim Glück, Hanna Preuss, Jeanne Droux, Upasana Maheshwari, Sucheta Sridhar, Susanne Wegener, Bruno Weber, Daniel Razansky, Mohammad El Amki, Andy Y. Shih, Annika Keller

## Abstract

Brain capillary pericytes are essential components of the neurovascular unit, yet the extent of their molecular heterogeneity within intact vascular networks remains poorly understood. Here, we combined spatial imaging with reanalysis of independent single-cell transcriptomic datasets to investigate the molecular organization of adult mouse brain pericytes. We identified spatial organization of pericyte molecular heterogeneity associated with anatomical region and position within the vascular network, including recurrent differences in ACE2, CASQ2, *Igf2,* and OPN expression. Moreover, pericyte molecular phenotypes varied with aging, acute ischemia, and circadian phase. Notably, light–dark phase emerged as a major axis of transcriptional variation, with pericytes exhibiting distinct circadian phase-associated molecular states. Together, these data demonstrate that adult brain pericytes exhibit spatially organized and dynamically regulated molecular heterogeneity associated with vascular and physiological context.

## Introduction

Pericytes are mural cells embedded within the vascular basement membrane of capillaries, where they contribute to blood–brain barrier (BBB) integrity, vascular stability, immune homeostasis, and cerebral blood flow regulation.^1–7^ Dysfunction of brain pericytes has been linked to aging, ischemic injury, and neurodegenerative disorders, conditions frequently associated with impaired neurovascular coupling, BBB dysfunction, and altered capillary perfusion.^3,8–12^While morphological and transcriptomic heterogeneity among brain pericytes is increasingly recognized^13,14^, the extent to which pericyte molecular phenotype varies across anatomical, vascular, and physiological contexts remains unclear.

Morphological studies, beginning with Zimmermann’s original description, show a gradual phenotypic continuum of mural cells along the vascular tree, extending from vascular smooth muscle cells (VSMCs) on arterioles, through transitional mural cells at arteriole–capillary transitions, to capillary pericytes. ^13,15^ These mural cells differ in morphology, vascular coverage, expression of contractile proteins, and vascular position, and some have been associated with different roles in vascular contractility and neurovascular coupling.^4,6,7^ Although capillary pericytes can be morphologically classified into thin-strand and mesh forms, these morphological classes show relatively limited electrophysiological differences. Earlier transcriptomic studies have generally resolved brain pericytes as a single molecular population.^13,16–18^

Recent single-cell and single-nucleus transcriptomic studies in mouse and human brain vasculature have revealed substantially greater molecular heterogeneity among pericytes than previously recognized.^14,17,19–21^. Molecularly distinct pericyte populations have been identified based on differential expression of signaling molecules, extracellular matrix proteins, transporters, and receptors.^14,19,21^ Additionally, brain mural cells originate from distinct embryonic sources depending on anatomical region, raising the possibility that developmental origin may contribute to adult pericyte diversity. Vascular transcriptomic studies have demonstrated pronounced zonation of endothelial gene expression along the arteriovenous axis of the brain vasculature^17,22–24^, and emerging evidence supports positional heterogeneity among pericytes.^20,21,25^ However, direct spatial mapping of molecularly distinct pericyte populations within intact brain vasculature remains limited, as reconstructing continuous arteriovenous topology across large capillary networks, together with pericyte molecular phenotype, is technically challenging.

Understanding the spatial organization of molecularly distinct pericytes is particularly important in aging and cerebrovascular disease, where pericyte alterations are often reported as global changes in pericyte abundance or vascular coverage.^11,12,26,27^ Consequently, changes affecting molecularly or spatially distinct pericyte populations may have been obscured when pericytes were analyzed as a single population. Furthermore, accumulating evidence indicates that vascular and perivascular cell states are influenced by anatomical, physiological, and pathological contexts^22–24,28^, suggesting that adult capillary pericyte phenotype may be dynamic rather than fixed.

Prompted by an initial observation of heterogeneous angiotensin-converting enzyme 2 (ACE2) expression in brain pericytes, we investigated how molecular heterogeneity among adult mouse brain capillary pericytes is spatially organized and whether it varies across physiological and pathological contexts. We show that subsets of neighboring pericytes differ in ACE2 expression independently of the canonical pericyte marker CD13 intensity, and use ACE2 expression levels to define ACE2*^high^* and ACE2*^low^* pericyte groups for spatial analysis. Three-dimensional reconstruction of intact cortical vascular networks revealed positional bias in ACE2 expression, with higher ACE2-associated vessel edges enriched near α-smooth muscle actin (ASMA)-positive arterioles. Reanalysis of independent single-cell (sc)RNA-seq datasets revealed additional dimensions of molecular heterogeneity among brain pericytes that extended beyond ACE2 expression, including differential expression of *Casq2*, *Spp1*, and *Igf2*, which we validated spatially in brain tissue. Distinct aspects of pericyte molecular heterogeneity also varied with aging, acute ischemia, and circadian phase. Collectively, these findings demonstrate that adult brain capillary pericytes exhibit region-, topology-, and context-dependent molecular heterogeneity.

## RESULTS

### Heterogeneity in ACE2 expression by pericytes in the mouse CNS

Angiotensin-converting enzyme 2 (ACE2) was recently identified as a marker of brain pericytes.^29^ Immunostaining for ACE2 demonstrated widespread expression of ACE2 among mural cells within the mouse CNS vasculature (Fig. S1A-C). ASMA-positive mural cells expressed ACE2, but at lower levels than capillary pericytes (Fig. S1A). In contrast, mural cells covering larger veins lacked detectable ACE2 expression (Fig. S1B). Nevertheless, elongated ACE2-positive pericyte processes frequently extended from adjacent capillary segments onto larger veins (Fig. S1B). ACE2 immunoreactivity was also observed in retinal mural cells, including both ASMA-positive ensheathing pericytes and adjacent ASMA-negative capillary pericytes (Fig. S1C). These observations are consistent with previous transcriptomic and morphological studies that describe gradual phenotypic transitions of mural cells along the arterio-venous axis, including a progressive reduction of contractile markers after the arteriole–capillary transition zones and poorly defined boundaries between capillary- and venous-associated pericytes.^13,17,20^

Within the capillary bed, neighboring pericytes frequently exhibited different levels of ACE2 expression, with some pericytes displaying weak or nearly undetectable ACE2 immunoreactivity (Fig. 1A, B), an observation not previously reported in the brain.^29,30^ Figure 1A shows two adjacent pericytes with differing ACE2 expression levels, where the variation in ACE2 immunoreactivity delineates the boundaries of neighboring pericyte territories.

**Figure 1.**
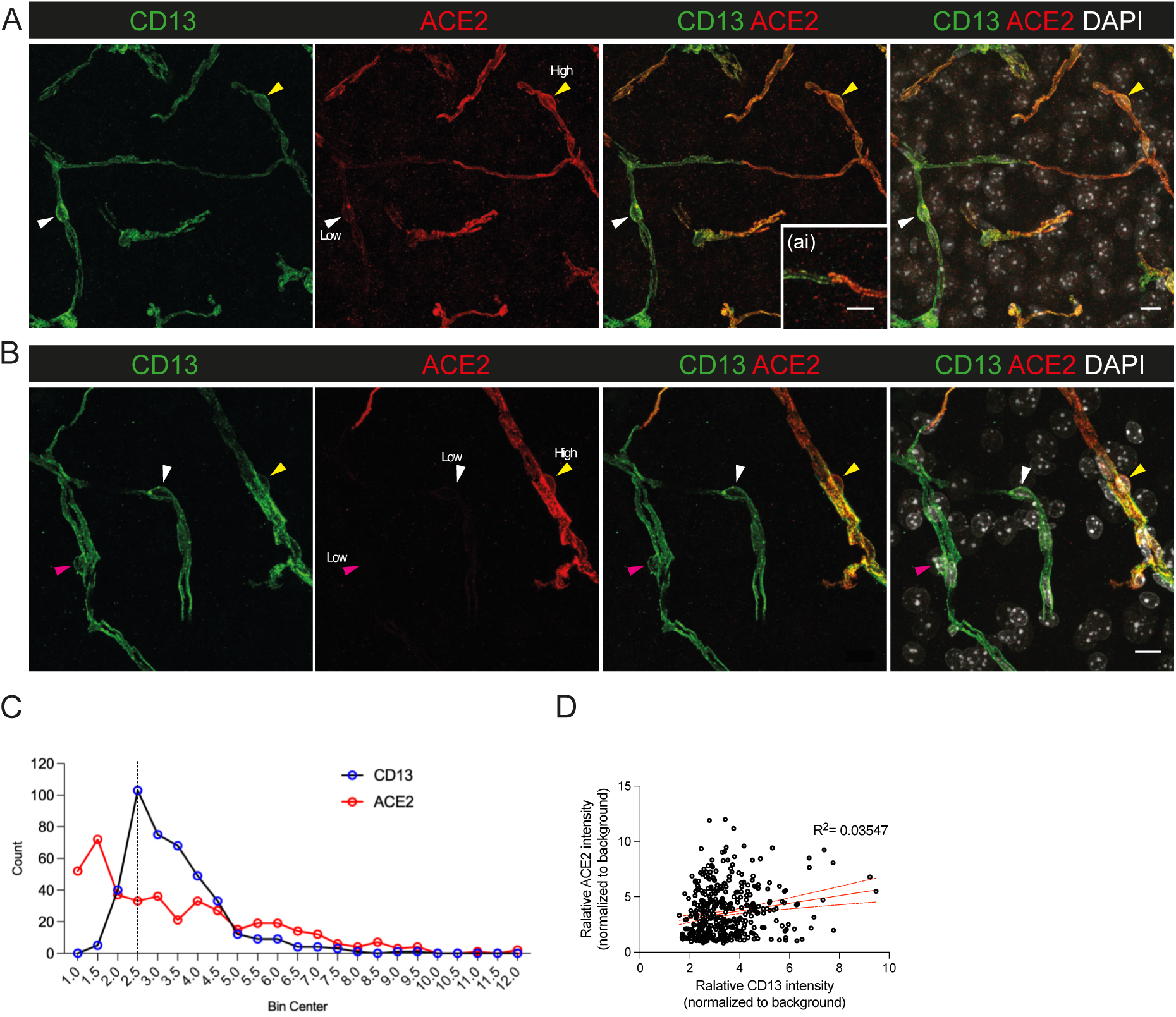
Heterogeneity in ACE2 expression by pericytes in the mouse CNS. **A** Variable ACE2 (in red) expression in hypothalamic pericytes in 2–3-month-old mouse brain. Adjacent ACE2*^low^* pericyte (white arrowhead) and ACE2*^high^* pericyte (yellow arrowhead) on the same capillary segment. **Ai** a higher magnification of the contact area between the processes of ACE2*^high^* and ACE2*^low^* pericyte. Cell nuclei are visualized with DAPI (in white), and pericytes are identified by CD13 (in green) immunostaining. Scale bars, 10 µm (**A**) and 2.5 µm (**Ai**). **B** Mesh and thin-strand pericytes express ACE2 (in red) variably. Arrowheads indicate an ACE2*^low^* thin-strand pericyte (white arrowhead), an ACE2*^low^* mesh pericyte (pink arrowhead) and an ACE2*^high^* mesh pericyte (yellow arrowhead). Cell nuclei are visualized with DAPI (in white), and pericytes are identified by CD13 (in green) immunostaining. Scale bar, 10 µm. **C** Frequency distribution of CD13 (blue) and ACE2 (red) immunofluorescence signal intensities in cortical pericytes. Intensities were binned at 0.5-unit intervals and plotted against bin centers. The dashed black vertical line denotes the threshold that defines the ACE2*^high^* and ACE2*^low^* pericyte populations. The pan-pericyte marker CD13 shows a unimodal distribution, whereas ACE2 displays a broader, complex distribution, reflecting expression heterogeneity. *n* = 418 pericytes examined from 3 independent mice. Confocal images are presented as maximum intensity projections. **D** The ACE2 expression level varies largely independently of CD13 expression level in pericytes. Each data point represents an individual cell; *n* = 418 pericytes examined from 3 mice. The red line shows the best-fit regression line, and the dashed red lines show the 95% confidence interval. r = 0.188, P < 0.001, R^2^ = 0.035, two-tailed Pearson correlation coefficient.

Quantification of CD13 and ACE2 signals in individual pericytes revealed a distinct distribution pattern for each marker (Fig. 1C). CD13 expression exhibited a unimodal distribution, indicating relatively uniform expression among pericytes. In contrast, ACE2 expression displayed a broader and more complex distribution, with values extending toward higher expression bins (Fig. 1C). Analysis of protein expression levels in individual cells showed no correlation between CD13 and ACE2 expression levels (Fig. 1D). Pericytes with differing ACE2 expression levels also demonstrated comparable PDGFRB immunoreactivity (Fig. S1D). Variability in ACE2 immunostaining intensity was also observed in capillary pericytes in the retina (Fig. S1E).

In summary, ACE2 expression reveals molecular heterogeneity among pericytes within local microvascular networks in the mouse CNS.

### Regional and vascular-topological organization of ACE2-defined capillary pericyte groups

To determine whether heterogeneity in pericyte ACE2 expression shows brain regional specificity, immunofluorescence-stained serial vibratome sections were first imaged using a fluorescence slide scanner, followed by targeted confocal imaging of selected anatomical regions (Fig. S2A). Images acquired from serial brain sections revealed anatomical differences in the abundance of lower-ACE2-expressing pericytes (Fig. 2A-C and Fig. S2B). Although ACE2 expression varied continuously among pericytes, a subset of cells exhibited relatively high ACE2 signal intensity (Fig. 1C). For subsequent spatial analyses, we therefore operationally classified pericytes into ACE2*^high^* and ACE2*^low^* (see Materials and Methods). Quantification of CD13-positive pericytes and their ACE2 expression levels across anatomical regions showed that ACE2*^high^* pericytes constituted the predominant group in all analyzed brain regions, with the relative proportions of ACE2*^high^* and ACE2*^low^* pericytes varying between regions. In the hippocampus, somatosensory cortex, and motor cortex, approximately 10% of pericytes were classified as ACE2*^low^*, compared to 30–40% in the hypothalamus and corpus callosum (Fig. 2C). Comparable regional distributions of ACE2*^high^* and ACE2*^low^* pericytes were also observed in genetically labeled pericytes in *Atp13a5*-tdTomato mice (Fig. S2C, D), supporting the robustness of the observed regional heterogeneity using CD13-based pericyte identification.

**Figure 2.**
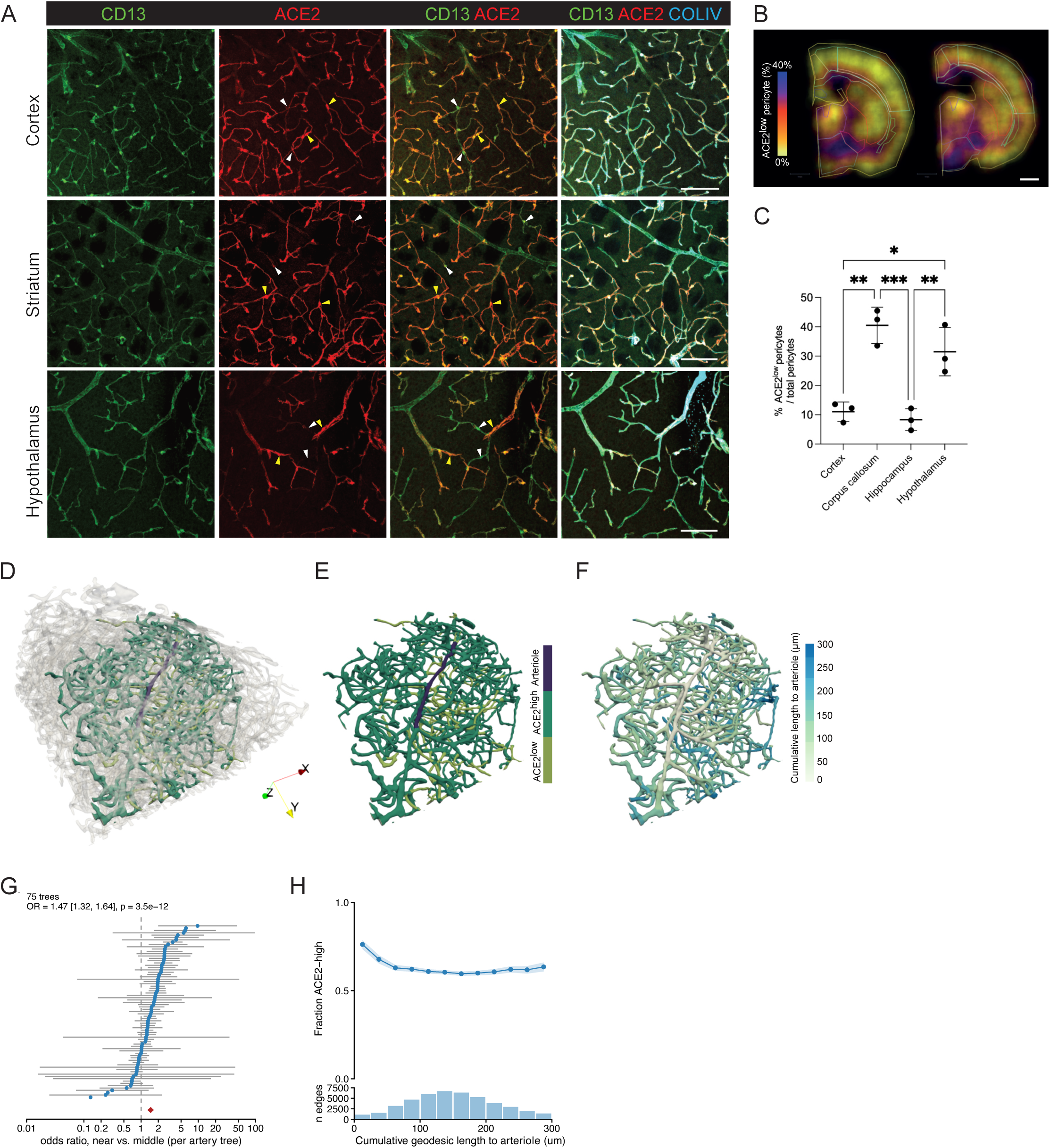
Regional and vascular-topological organization of ACE2-defined capillary pericyte groups. **A** The distribution of ACE2*^high^* (yellow arrowheads) and ACE2*^low^* (white arrowheads) pericytes in the cortex, striatum, and hypothalamus. ACE2 in red, pan-pericyte marker CD13 in green, and vascular basement membrane (COLIV) in blue. Scale bars, 100 µm. Confocal images are presented as maximum intensity projections. **B** Heatmap visualization of ACE2*^low^* pericyte proportions (%) across coronal brain sections. The color scale ranges from yellow (low proportion) to blue/purple (high proportion). Distinct regional heterogeneity in ACE2*^low^* pericyte abundance is evident between cortical and subcortical structures. Scale bar, 1 mm. **C** Quantification of the percentage of ACE2*^low^* pericytes relative to total pericytes across cortex, corpus callosum, hippocampus, and hypothalamus. Significant regional differences in ACE2*^low^* abundance are observed between gray matter structures and white matter/subcortical regions. Data are presented as mean ± SD. Each dot represents an individual animal (*n* = 3). \**p*<0.05, \*\**p*<0.01, \*\*\**p*<0.001, one-way ANOVA with Tukey’s multiple comparison. **D** Three-dimensional reconstruction of a representative cortical vascular volume showing an extracted vascular tree (green) within the surrounding vascular network (grey). **E** Extracted vascular tree showing ACE2*^low^* (light green) and ACE2*^high^* (dark green) vessel edges relative to an ASMA-positive arteriole (dark blue). **F** Cumulative geodesic length of vessel edges from the arteriole, visualized by color intensity (white, proximal; blue, distal). **G** Forest plot showing odds ratios (ORs, near vs. middle segments) for ACE2*^high^* edge localization across 75 individual vascular trees. Blue circles and horizontal grey bars denote individual ORs and 95% confidence intervals (CIs), respectively. The red diamond indicates the overall pooled OR (1.47, 95% CI 1.32–1.64, *P* = 3.5 × 10^⁻12^, *n* = 75 arteriolar territories). Between-territory heterogeneity was moderate to substantial (three-band model: I^2^ = 64%, Cochran’s Q *P* = 1.1 × 10⁻^14^; five-band model: I² = 49%, Cochran’s Q *P* = 2.8 × 10^⁻6^). **H** Fraction of ACE2*^high^* vessel edges as a function of cumulative geodesic length from the arteriole. The upper plot shows the mean fraction within each distance bin, with shading indicating the 95% confidence interval. The lower histogram shows the total number of vessel edges within each corresponding distance bin.

We next asked whether ACE2-defined pericyte groups were associated with distinct vascular territories. Previous studies have shown that mesh pericytes cover slightly larger capillaries than thin-strand pericytes in the mouse cerebral cortex.^13^ Because mesh and thin-strand pericytes exhibit heterogeneous ACE2 expression (Fig. 1B and Fig. S1E), we examined whether ACE2 expression was related to capillary diameter. However, no correlation was observed between pericyte ACE2 expression level and capillary diameter in the three analyzed brain regions (Fig. S2E-G). We therefore asked whether ACE2 expression was instead associated with vascular architecture. Cortical microvascular networks were reconstructed from ASMA-positive arterioles into the downstream capillary network using large-volume light-sheet imaging and simulation-based vessel segmentation (Fig. 2D and Fig. S2H). To assess the relationship between ACE2 expression and vascular position, ACE2 expression was analyzed at the level of vessel edges rather than individual pericyte somata, because pericytes at branch points could not be unambiguously assigned to a single vessel segment. In total, 44,924 vessel edges from 75 vascular trees were included in the positional analysis. An extracted vascular tree illustrates that ACE2*^high^* capillaries are predominantly localized within the proximal capillary network (Fig 2E-F). This spatial distribution was further validated by confocal imaging, which revealed the same pattern (Fig S2I). Quantification of cumulative geodesic distance from the nearest arteriole revealed that ACE2*^high^* capillary edges were most frequent near ASMA-positive arterial termini and progressively decreased along the proximal capillary network before reaching a lower plateau in more distal segments (Fig. 2G-H). Accordingly, ACE2*^high^* capillaries were significantly enriched in proximal compared with more distal capillary segments. This positional bias remained significant across multiple analytical approaches, including different distance-binning strategies and complementary branch-order analysis (Fig. S2J-K). Together, these findings indicate that ACE2 expression exhibits a positional bias along the microvascular tree, with higher expression in ASMA-negative pericytes proximal to arterioles, while remaining heterogeneously distributed throughout the distal capillary bed.

Collectively, these findings indicate that ACE2-defined capillary pericyte heterogeneity is spatially organized by anatomical region and shows additional positional bias within the cortical vascular network.

### Transcriptional heterogeneity of mouse brain pericytes

The regional and vascular-topological heterogeneity among ACE2-defined pericyte populations prompted us to examine whether molecular heterogeneity in mouse brain pericytes could also be detected in published transcriptomic datasets. In prior analyses of the Vanlandewijck et al.^17^ and Yao et al.^31^ datasets, molecular heterogeneity within capillary pericytes was not specifically examined. In contrast, Smith et al.^19^ identified eight transcriptionally distinct hypothalamic pericyte subclusters, although this predicted heterogeneity was not validated in tissue. To address these gaps, we first reanalyzed a publicly available scRNA-seq dataset generated by Vanlandewijck et al., followed by independent analyses of the hypothalamus-focused dataset from Smith et al. and the whole-brain dataset from Yao et al. (S.Table 1). Clustering of the Vanlandewijck dataset identified three endothelial clusters (capillary (capEC), arterial (aEC), and venous (vEC) endothelial cells), together with perivascular macrophages (PVM), perivascular fibroblasts (PVF), and four mural cell clusters, comprising two smooth muscle cell clusters (SMC1, SMC2) and two pericyte (Ace2_high PC, Ace2_low PC) clusters (Fig. 3A and Fig. S3A). Despite the relatively limited number of pericytes captured in this dataset (1005 cells) (S.Table 1), unsupervised clustering identified two transcriptionally distinct pericyte populations. One cluster (Ace2_high PC, 530 pericytes) was characterized by enriched expression of *Ace2, Casq2, and Ednrb*, with *Ace2* differential expression consistent with the heterogeneous ACE2 expression observed in situ. The second cluster (Ace2_low PC, 475 pericytes) showed enriched expression of *Igf2*, *Spp1*, and *Slc1a3*, among other genes (Fig. 3A, B; Fig. S3A, B and S.Table 1, 2). These genes (*Igf2*, *Spp1*, *Slc1a3*) are also expressed by PVF; however, cells in the Ace2_low PC cluster retained canonical pericyte markers (e.g., *Anpep*, *Kcnj8*, *Abcc9*, *Pdgfrb*, *Vtn*) and lacked PVF markers (e.g., *Pdgfra*, *Cemip*, *Col1a1*) (Fig. S3C), confirming their pericyte identity. CellChat (Fig. S3D-G) and NATMI (S.Table 2) analyses suggested differences in predicted signaling interactions between ACE2_high PC and ACE2_low PC, and neighboring capillary endothelial cells.

**Figure 3.**
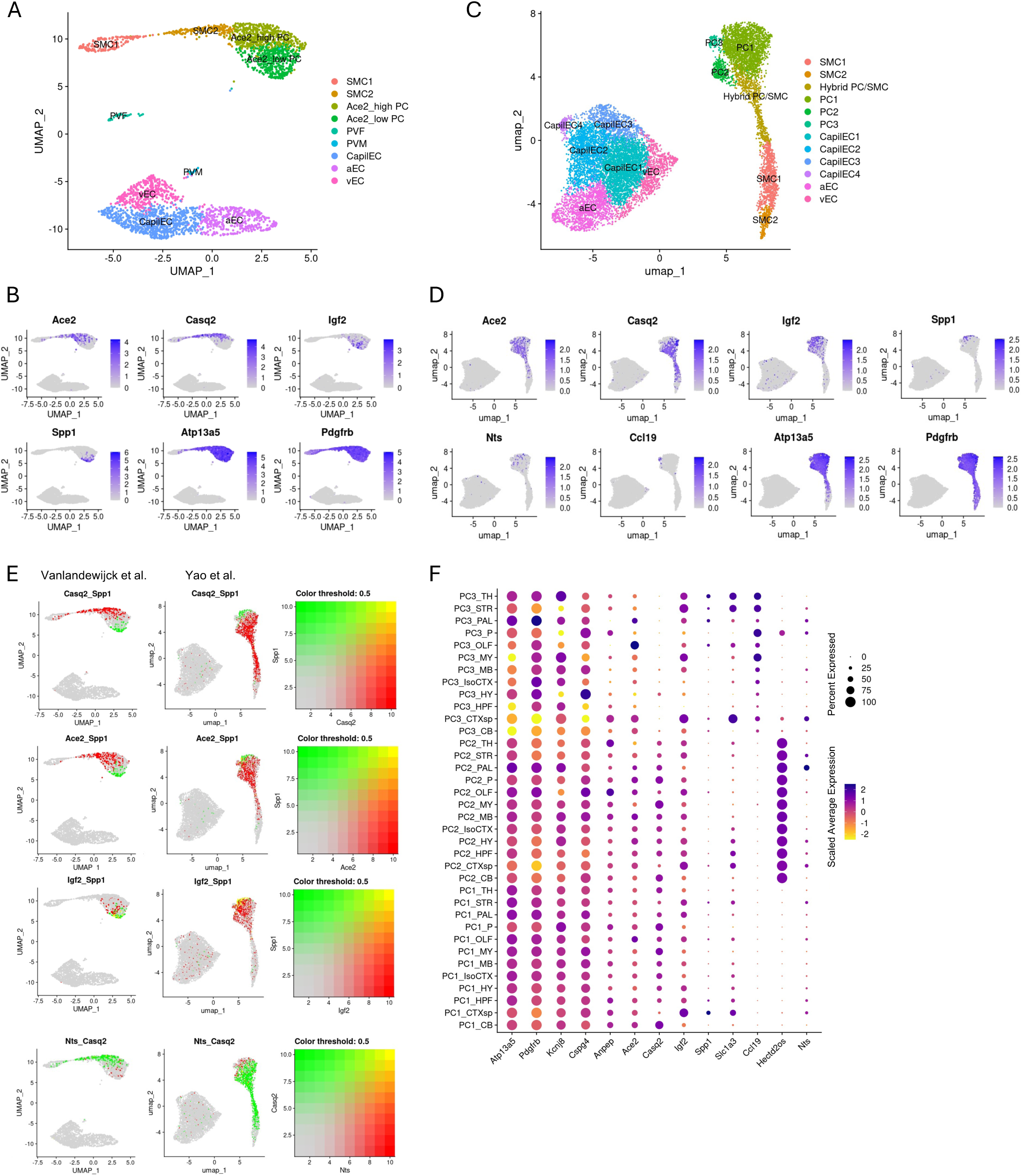
Transcriptional heterogeneity of mouse brain pericytes. **A** UMAP projection of cell clusters from the Vanlandewijck et al. dataset^17^ at a resolution of 0.8 and colored according to cell clusters. **B** Feature plots displaying expression profiles of cluster-specific (*Ace2, Casq2, Igf2, Spp1*) and pan-pericyte markers (*Atp13a5*, *Pdgfrb*) within the Vanlandewijck et al.^17^ dataset. **C** UMAP projection of endothelial and mural cell clusters from the Yao et al. dataset^31^ at a resolution of 0.8 and colored according to cell clusters. **D** Feature plots displaying expression profiles of selected genes (*Ace2, Casq2, Igf2, Spp1, Nts, Ccl19*) and pan-pericyte markers (*Atp13a5*, *Pdgfrb*) within the mural cell cluster from the Yao et al. dataset^31^. **E** Bivariate feature plots highlighting single-cell co-expression of a pan-pericyte marker alongside selected pericyte markers. Green and red signal intensities represent single-gene expression, while yellow indicates co-expression. **F** Dot plots showing selected marker expression across integrated pericyte subclusters from various brain regions in the Yao et al. dataset^31^. Dot size represents the percentage of cells expressing the target gene, and color intensity reflects scaled average expression level (purple, high; yellow, low). CB, cerebellum; CTX, cerebral cortex; CTXsp, cortical subplate; HPF, hippocampal formation; HY, hypothalamus; IsoCTX, isocortex; MB, midbrain; OLF, olfactory areas; P, pons; PAL, pallidum; STR, striatum; TH, thalamus.

We next reanalyzed vascular cells from a published scRNA-seq dataset generated from *Gipr-*expressing cells isolated from the adult mouse hypothalamus^19^. Because relatively few female vascular cells were recovered, and female and male vascular cells clustered separately (Fig. S4A), subsequent analyses were performed using data from male mice, which yielded the highest cell count. Unbiased clustering of endothelial and mural cells resulted in eight clusters: one endothelial cell cluster (EC), two smooth muscle cell clusters (SMC1, SMC2), and five pericyte clusters (PC1-5, in total 2615 cells) (Fig. S4B and S.Table 1, 3). *Ace2* was not differentially expressed across pericyte clusters; however, as seen in the ACE2_low cluster in the Vanlandewijck dataset (Fig. 3A, B), one pericyte cluster (PC1) expressed higher levels of *Spp1* and *Gatm* and lower levels of *Casq2* (Fig. S4C). PC2 expressed higher levels of *Mgp* and *Serpini1*, and PC3 was enriched for *Kcnj8*, *Abcc9*, and *Casq2*. PC4 expressed higher levels of *Slc12a2*, *Lars2, Notch3, Lama2, Casq2* (Fig. S4C). PC5 (approx. 8 % of pericytes) displayed high *Nts* expression (Fig. S4C, D). Together, these expression patterns indicate molecular heterogeneity among hypothalamic pericytes that extends beyond a single marker or binary classification.

We further re-analyzed the endothelial and mural cells from the scRNA-seq dataset generated by Yao et al.^31^, which contains both female and male cells across 12 brain regions. In this dataset, we identified six endothelial cell clusters: venous (vEC), arterial (aEC), and four capillary (capEC 1-4). In addition to two smooth muscle cell clusters (SMC1, SMC2), we identified three pericyte clusters (PC 1-3, 2474 cells) and one mural cell cluster displaying features of both pericytes and smooth muscle cells (hybrid PC/SMC; 669 cells, approximately 16% of mural cells) (Fig. 3C; Fig. S5A and S.Table 1, 4). Two smaller pericyte clusters, PC3 (approx. 4% of pericytes) and PC2 (approx. 11% of pericytes), were defined by the expression of Ccl19 and the long non-coding RNA *Hectd2os*, respectively. Although the major pericyte cluster PC1 (2089 cells) did not further separate into stable subclusters at higher clustering resolution, differential expression of several genes identified in the Vanlandewijck and Yao datasets, including *Ace2*, *Casq2*, *Igf2*, *Spp1*, and *Nts*, remained evident among individual pericytes (Fig. 3E). There were no apparent sex-specific differences; all clusters contained cells from both male and female mice (Fig. S5B, C). Similarly, PC1-PC3 and the hybrid PC/SMC cluster were distributed across anatomical regions without clear region-specific enrichment (Fig. 3F and Fig. S5D). Nevertheless, expression levels of several genes, including *Igf2* and *Spp1*, varied across anatomical regions, with the highest expression observed in pericytes isolated from the cortical subplate (Fig. 3F).

Although the three analyzed datasets differed in their cell isolation approaches, anatomical regions, and scRNA sequencing methodologies, they all contained pericytes that showed largely mutually exclusive expression patterns for *Ace2* and *Spp1*, and for *Casq2* and *Spp1* (Fig. 3B, D, E and Fig. S4D, E, F). However, the co-expression pattern between *Igf2* and *Spp1* differs between datasets: the Vanladerwijck et al. dataset primarily suggests a mutually exclusive expression pattern between *Igf2* and *Spp1,* but the Yao dataset reveals a substantial subpopulation of cells co-expressing both *Igf2* and *Spp1*, alongside cells expressing only one or the other (Fig. 3E). The Smith et al. dataset, originating from the hypothalamus, contains a cluster of strongly *Nts-*positive pericytes (cluster PC5) that shows a largely mutually exclusive expression pattern with *Casq2* (Fig. S4F). Although *Nts*-expressing pericytes do not form a separate cluster in the Vanladerwijck and Yao datasets, these *Nts*-positive pericytes in those datasets also show a largely mutually exclusive expression pattern with *Casq2* (Fig. 3E). Importantly, pericytes across all analyzed datasets consistently showed widespread co-expression of canonical pericyte markers such as *Pdgfrb*, *Atp13a5*, *Abcc9,* and *Kcnj8*, confirming their core pericyte identity (Fig. S5E).

In summary, reanalysis of three independent single-cell transcriptomic datasets showed recurrent yet context-dependent molecular heterogeneity among adult mouse brain pericytes. Marker expression patterns partially overlapped across datasets, supporting the presence of multiple molecularly distinct pericyte groups.

### Convergent transcriptional regulatory programs in brain mural cells

Brain mural cells have diverse developmental origins: those in the telencephalon are derived from neural crest, hindbrain mural cells originate from the mesenchyme, and diencephalon mural cells have a mixed origin.^32–35^ We therefore asked whether differences in developmental origin could contribute to the molecular heterogeneity observed among adult brain pericytes, or whether mural cells of distinct origins converge on similar regulatory programs. Interestingly, transcription factor (TF) activity analysis of both VSMCs and pericytes from these different brain regions revealed that they clustered by cell type rather than by region (Fig. S6A). Furthermore, analysis of TF activity, including comparisons with endothelial cells, revealed that both VSMCs and pericytes share a common TF activity signature (Fig. S6B), characterized by transcription factors such as MSX1 and FOXF1, which are linked to the development of mesoderm-derived structures and the vasculature^6^. This suggests that brain mural cells, despite their diverse embryonic origins, converge on similar regulatory programs in the adult brain. A clear divergence in TF activity was nevertheless observed between pericytes and VSMCs (Fig. S6A), indicating that this convergence preserves distinct regulatory identities for each mural cell class.

### Spatial validation of molecular heterogeneity among brain pericytes

Based on our analysis of three independent scRNA-seq datasets, we identified several genes, including *Igf2*, *Spp1*, and *Casq2*, that were expressed heterogeneously among brain pericytes (Fig. 3 and Fig. S3-S5). To determine whether this transcriptional heterogeneity could also be detected within intact brain vasculature, we performed in situ hybridization and immunohistochemistry.

A recent study demonstrated that the pericyte-derived IGF2 in the hippocampus plays a role in long-term memory formation^36^. Our transcriptomic analyses suggested that *Igf2* is expressed by pericytes across multiple brain regions, although expression levels varied among individual pericytes (Fig. 3B, D, and Fig. S4D). In situ hybridization confirmed heterogeneous *Igf2* expression in cortical pericytes (Fig. 4A), with similar variability observed in the hippocampus and hypothalamus (Fig. S7A).

**Figure 4.**
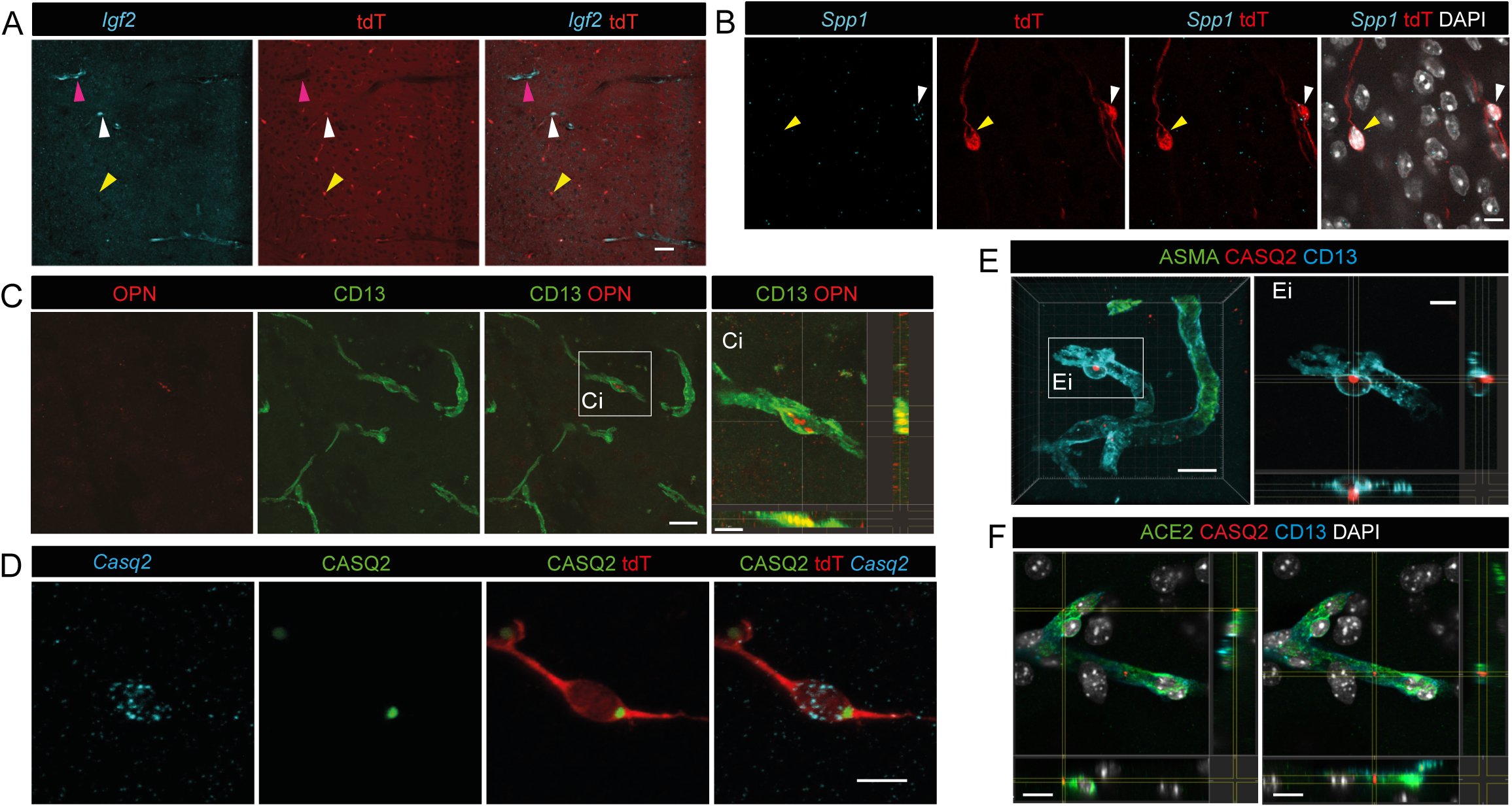
Spatial validation of molecular heterogeneity of brain pericytes. **A** *Igf2* mRNA expression (in cyan) in cortical pericytes (tdT, in red) and perivascular fibroblasts (pink arrowhead) within the cerebral cortex of an *Atp13a5*Cre-tdTomato mouse. White and yellow arrowheads indicate *Igf2*-positive and *Igf2*-negative pericytes, respectively. Scale bar, 100 µm. **B** *Spp1* mRNA expression (in cyan) in pericytes (tdT, in red) in the *Atp13a5*Cre-tdTomato mouse brain. White and yellow arrowheads indicate *Spp1*-positive and *Spp1*-negative pericytes, respectively. Cell nuclei are visualized with DAPI (in white). Scale bar, 5 µm. **C** Osteopontin (OPN, in red) expression in pericytes (CD13, in green). **Ci** A magnified view with orthogonal projections (XZ and YZ planes) showing the intracellular localization of OPN. Scale bars, 10 µm (**C**) and 7 µm (**Ci**). **D** CASQ2 mRNA (in cyan) and protein (in green) expression in pericytes (tdT, in red) of *Atp13a5*Cre-tdTomato mouse brain. Scale bar, 10 µm. **E** CASQ2 (in red) immunoreactivity in pericytes (CD13, in cyan). ASMA (in green) labels VSMCs. **Ei** Magnified inset with orthogonal projections (XZ and YZ planes) showing CASQ2 localized as a single punctum within the pericyte cell body. Scale bars, 10 µm (E), 5 µm (Ei). **F** Images with orthogonal projections (XZ and YZ planes) showing CASQ2 (in red) localized as puncta along pericyte processes. Pericytes are labeled with CD13 (in cyan) and ACE2 (in green). Cell nuclei are visualized with DAPI (in white). Scale bars, 10 µm. All confocal images are presented as maximum intensity projections.

*Spp1* expression was detected in a relatively small subset of pericytes across all analyzed datasets (Fig. 3B, D, and Fig. S4D). Consistent with these findings, *Spp1* mRNA and its protein product, osteopontin, were detected only in a subset of pericytes in brain tissue (Fig. 4B, C and Fig. S7B, C). Osteopontin was also detected in PVF (Fig. S7C), consistent with the transcriptomic analyses showing *Spp1* expression in both pericytes and PVFs (Fig. S3C).

*Casq2*, which encodes the calcium-buffering protein calsequestrin 2 (CASQ2), has not previously been reported to be expressed in brain pericytes. We confirmed the expected *Casq2*/CASQ2 expression in cardiac myocytes and Purkinje cells (Fig. S7D) and further identified both *Casq2* mRNA and CASQ2 protein expression in brain pericytes (Fig. 4D). CASQ2 expression levels and subcellular localization varied among pericytes. CASQ2 immunoreactivity was detected both in the pericyte cell body and along processes (Fig. 4D-F), whereas *Casq2* mRNA was distributed more broadly throughout the cell body (Fig. 4D).

Taken together, these findings demonstrate that markers identified by transcriptomic analyses exhibit heterogeneous spatial expression patterns within the brain vasculature, supporting molecular heterogeneity among adult brain pericytes at both transcript and protein levels.

### Region-specific changes in the abundance of ACE2-defined pericyte groups and CASQ2 organization in cortical pericytes during aging

Having established that ACE2-defined pericyte groups exhibit regional and vascular-topological organization in young adult mice, we next investigated whether their relative abundance changes with age. We compared the proportions of ACE2*^high^* and ACE2*^low^* pericytes between young adult and 14–17-month-old mice across selected brain regions. Immunofluorescence revealed an increased proportion of ACE2*^low^* pericytes in the somatosensory cortex and hippocampus of aged mice compared to young adults (Fig. 5A, S8). In deep brain regions, where the proportion of ACE2*^low^* pericytes was already higher in young adults (e.g., thalamus), no further increase was observed. In young mice, ACE2*^low^* pericytes in the somatosensory cortex were predominantly localized to cortical layer VI; however, in aged mice, the proportion of ACE2*^low^* pericytes was increased across all cortical layers (Fig. 5B). Cortical pericyte numbers were not reduced in aged mice compared to young adults (Fig. 5C). These results suggest that the increased abundance of ACE2*^low^* pericytes in the aged cortex reflects phenotypic remodeling rather than the selective loss of ACE2*^high^* pericytes, and that this remodeling is accompanied by a wider spatial distribution of molecularly heterogeneous pericytes across cortical layers during aging.

**Figure 5.**
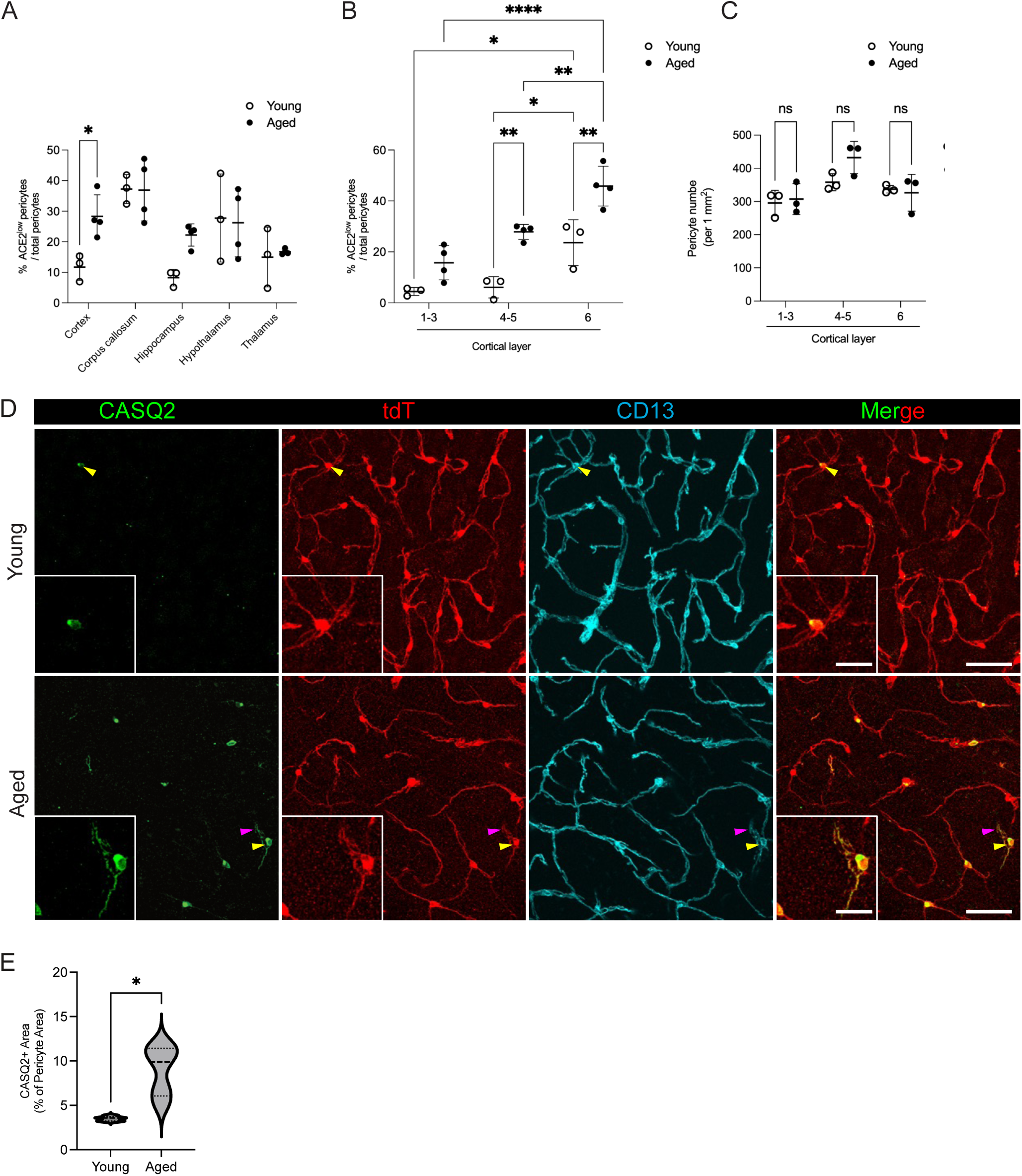
Region-specific changes in the abundance of ACE2-defined pericyte subtypes and CASQ2 immunoreactivity in aged mice. **A-B** Quantification of the proportion of ACE2*^low^* pericytes across different brain regions (**A**) and layers of the cerebral cortex (**B**) in young (2–3-month-old) and aged (14–17-month-old) mice, determined by CD13 and ACE2 immunostaining. *n* = 3–4 mice. **C** Quantification of pericyte numbers in young and aged animals across cortical layers. *n* = 3–4 mice. Data are presented as mean ± SD. \**p* <0.05, \*\**p*<0.01, \*\*\*\**p*<0.0001, one-way ANOVA (A, C) and 2-way ANOVA (B), followed by Tukey’s multiple comparison test. **D** Representative confocal images illustrating reduced CASQ2 (in green) immunoreactivity and altered intracellular punctate localization in cortical pericytes (labeled by tdT and CD13, in red and cyan, respectively) from young (2-3 months old) and aged (14-17 months old) mice. Yellow arrowheads indicate CASQ2-positive pericytes. Purple arrowheads indicate CASQ2 puncta on pericyte processes. Indicated pericytes are shown at higher magnification in the insets. Scale bars, 50 µm (main images) and 20 µm (insets). Confocal images are presented as maximum intensity projections. **E** Quantification of the intracellular CASQ2-positive area within CASQ2-positive cortical pericytes. The thick dotted lines represent the median. The thinner dotted lines show the interquartile range from the 25^th^ to the 75^th^ percentile. Data are presented as mean ± SD. \**p*<0.05. Unpaired *t*-test with Welch’s correction. *n* = 5 mice per group (268 cells from the young animals and 370 cells from the aged animals). Unpaired *t*-test with Welch’s correction. *p* = 0.0116.

Beyond altered ACE2 expression patterns, pericytes in the somatomotor cortex of aged mice showed increased CASQ2 immunoreactivity and larger intracellular CASQ2 puncta within the cell soma (Fig. 5D). Quantification of the CASQ2-positive area per cell confirmed a significant increase in CASQ2-positive intracellular area (Fig. 5E).

Together, these findings indicate that aging is associated with region-dependent remodeling of pericyte molecular phenotype, including altered proportions of ACE2-defined pericyte groups and changes in CASQ2 immunoreactivity.

### Acute ischemia alters CASQ2 organization in cortical pericytes

The aging-associated remodeling of pericyte molecular phenotype prompted us to investigate whether similar changes could also occur during the acute phase of ischemic stroke. To address this, we performed thrombin-induced occlusion of the middle cerebral artery (MCAO) and analyzed pericyte molecular phenotypes 4 h after ischemia induction (Fig S9A). Quantification of CD13 immunoreactivity revealed no significant difference between sham-operated and ischemic mice (Fig. S9B, C), indicating that the canonical pericyte marker remained stable during the early post-ischemic phase. We next examined the distribution of ACE2-defined pericyte groups. Within the ipsilateral hemisphere, the proportions of ACE2*^high^* and ACE2*^low^* pericytes in the ischemia-affected somatosensory cortex and in the hypothalamus did not differ between sham-operated and MCAO mice (Fig. S9B, D), indicating that the relative abundance of ACE2-defined pericyte groups was preserved at this early time point after ischemia.

In contrast, the CASQ2 immunostaining pattern in pericytes was markedly altered following ischemia. In sham-operated mice and in the contralateral hemisphere, CASQ2 immunoreactivity in cortical pericytes was predominantly restricted to discrete puncta (Fig. 6A), similar to the pattern observed in naïve mice (Fig. 4E, F). In the ischemic core, however, CASQ2 immunoreactivity in pericytes frequently became diffuse, extending throughout the cell body and processes (Fig. 6A). Quantification of CASQ2-positive signal area within individual pericytes confirmed a significant increase in intracellular CASQ2 occupancy following ischemia (Fig. 6B).

**Figure 6.**
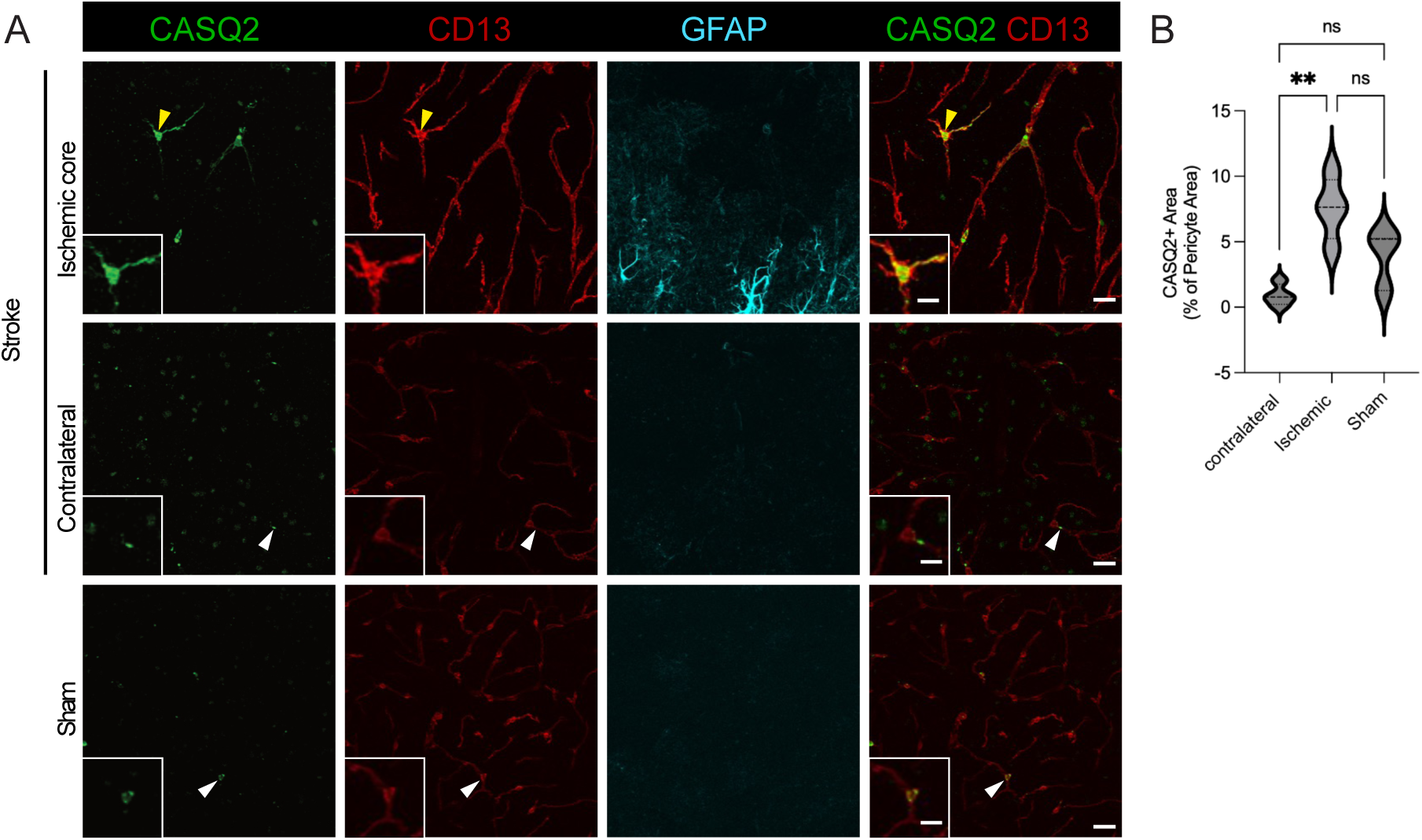
Acute ischemia alters CASQ2 organization in cortical pericytes. **A** Representative confocal images showing altered CASQ2 immunoreactivity (in green) in cortical pericytes identified by CD13 (in red) 4 h after middle cerebral artery occlusion (MCAO). In the ischemic hemisphere, CASQ2 frequently exhibited a diffuse intracellular distribution in pericytes (yellow arrowheads), whereas CASQ2 immunoreactivity remained predominantly punctate in the contralateral hemisphere and in sham animals (white arrowheads). Indicated pericytes are shown at higher magnification in the insets. GFAP (in cyan) labels reactive astrocytes within the ischemic core. Scale bars, 20 µm (main images) and 10 µm (insets). Confocal images are presented as maximum intensity projections. **B** Quantification of the intracellular CASQ2-positive area within cortical pericytes. *n* = 4 per group. The thick dotted lines represent the median. The thinner dotted lines show the interquartile range from the 25th to the 75th percentile. Data are presented as mean ± SD. \*\**p*<0.01; ns, not significant. One-way ANOVA followed by Tukey’s multiple-comparison test.

These findings indicate that acute ischemia rapidly remodels the intracellular organization of CASQ2 in pericytes, whereas the distribution of ACE2-defined pericyte groups remains unchanged during the early post-ischemic period.

### Circadian phase defines a major axis of transcriptional heterogeneity in brain pericytes

Having found that pericyte molecular features can change acutely following ischemia, we next asked whether pericyte heterogeneity also varies across normal physiological states. The analyses presented above for the Yao et al. dataset were restricted to cells collected during the light phase of the circadian cycle (Fig. 3C–F, Fig. S5). Because the Yao et al. dataset also included cells collected during the dark phase, we used these transcriptomes to test whether circadian phase contributes to pericyte transcriptional heterogeneity. Incorporation of dark-phase transcriptomes revealed a clear separation of pericytes by circadian phase in UMAP space, a pattern not observed in endothelial cells or VSMCs (Fig. 7A, B). Unbiased clustering of mural cells (7641 cells) identified four VSMC clusters (SMC 1-4), four pericyte clusters (PC 1-4), and one hybrid PC/SMC cluster (Fig. 7C, Fig. S10A, S.Table 4). Two major pericyte transcriptional groups (PC1 and PC2) showed a strong association of cluster membership with circadian phase (χ^2^=2382.45, *df*=1, *P*<0.0001, χπ=0.81): one population (PC1, 1846 of 2026 cells) was predominantly composed of light-phase cells, and the other (PC2, 1447 of 1609 cells) was enriched for dark-phase cells (Fig. 7D, STable 4). In contrast, no sex-associated separation was detected (Fig. S10B). Although the dataset included pericytes from multiple brain regions, assessment of whether phase-associated transcriptional programs differ between regions would require balanced region-by-phase sampling, which was not available in this dataset (S.Table 4, Fig. S10C).

**Figure 7.**
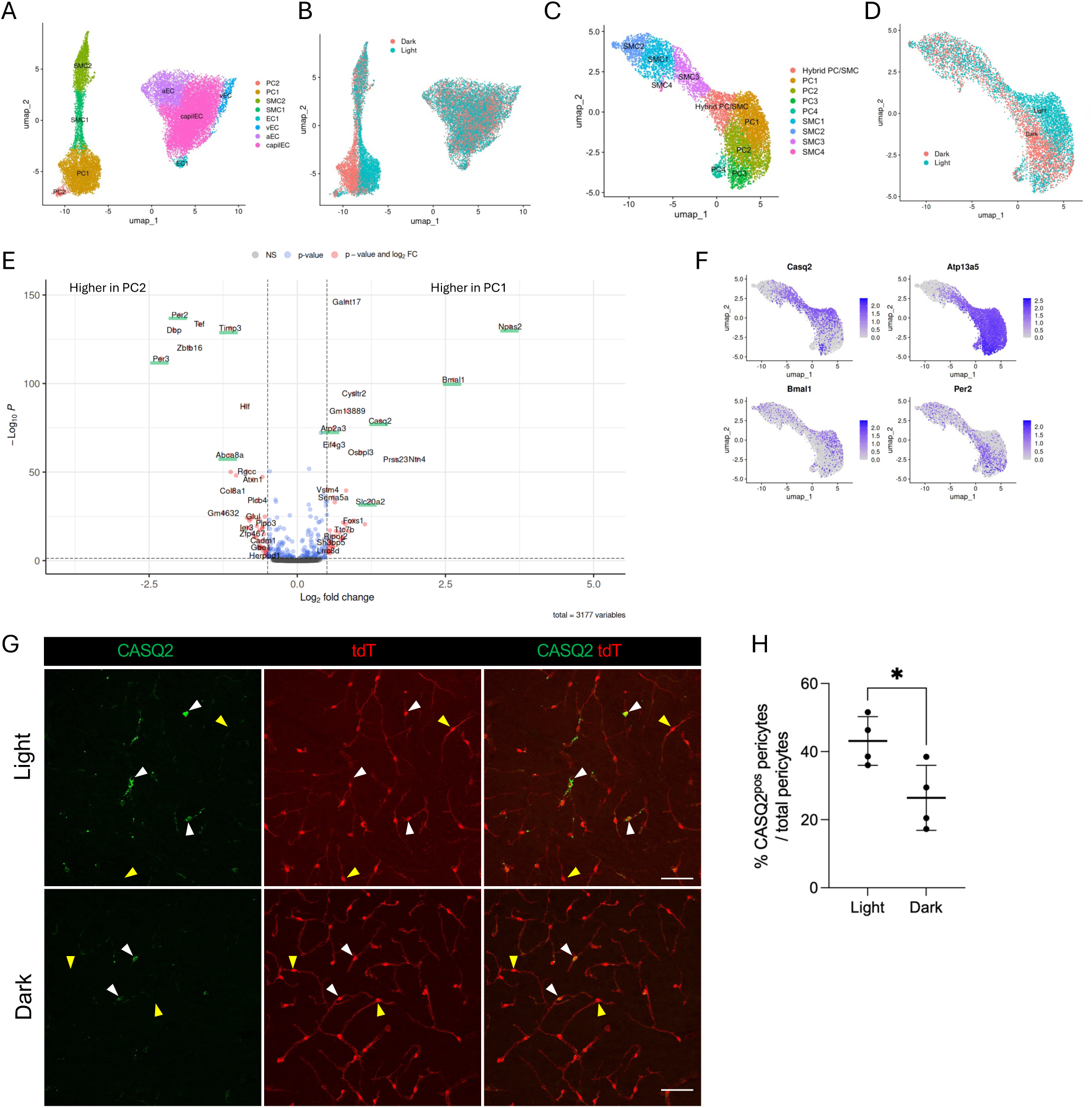
Circadian phase defines a major axis of transcriptional heterogeneity in brain pericytes. **A** UMAP visualization of endothelial and mural cell populations from the Yao et al. dataset, including cells collected during the light and dark phases. **B** UMAP showing endothelial and mural cells labeled according to circadian phase (light and dark) at the time of tissue collection. (Annotated in the original dataset as “medical_conditions”). **C** UMAP visualization of identified mural cell clusters. **D** UMAP of mural cells labeled according to circadian phase (light and dark) at the time of tissue collection. **E** Volcano plot showing the differential gene expression between two pericyte subclusters (PC1 and PC2). Each dot represents an individual gene plotted by log_2_ fold change (x-axis) and −log_10_ adjusted *p-*value (y-axis). Differentially expressed genes are in red. Dashed lines indicate the significance thresholds (adjusted *p*-value<0.05 and |log_2_FC|>0.25). Selected genes of interest are underlined in green. **F** Feature plots showing the expression of pericyte genes (*Casq2*, *Atp13a5*) and circadian-related genes (*Bmal1* and *Per2*) within mural cells. **G** CASQ2 (in green) expression in cortical pericytes (tdT, in red) during the light phase and dark phase in *Atp13a5*Cre-tdTomato mice. White arrowheads indicate the CASQ2-positive and yellow arrowheads indicate CASQ2-negative pericytes. Scale bars, 50 µm. Confocal images are presented as maximum intensity projections. **H** Quantification of the proportion of CASQ2-positive pericytes in samples across the isocortex during light and dark phases. Each data point represents one animal (*n* = 4). Data are presented as mean ± SD. \**p*<0.05. Unpaired *t*-test with Welch’s correction.

Pseudo-bulk RNA-seq analysis comparing PC1 and PC2 clusters revealed a coordinated light–dark transcriptional program rather than differential expression of individual clock genes alone. Dark-phase pericytes were enriched for canonical circadian output genes, including *Per2*, *Per3*, *Dbp*, *Tef*, and *Hlf*, as well as genes linked to metabolic, extracellular matrix, and stress-response pathways, such as *Glul*, *Timp3*, *Abca8a*, and *Ier3*. In contrast, PC1 pericytes showed higher expression of *Bmal1/Arntl* and *Npas2*, accompanied by the expression of genes associated with calcium and ion handling, membrane transport, and extracellular or secretory regulation, including *Casq2*, *Atp2a3*, *Slc20a2*, *Galnt17*, *Ntn4*, and *Prss23* (Fig. 7E and Fig. S10D). Transcription factor activity analysis further identified distinct regulatory signatures associated with circadian clock programs (Fig. S10E).

*Casq2* was among the genes enriched in light-phase-associated pericytes (Fig. 7E, F, and Fig. S10D). Consistent with the transcriptomic analysis, CASQ2 immunoreactivity in cortical pericytes also exhibited a circadian phase-dependent pattern, with a higher proportion of CASQ2-positive pericytes observed during the light phase (Fig. 7G, H). In contrast, immunoreactivity for the pan-pericyte markers (CD13 and PDGFRB), the heterogeneously expressed pericyte marker ACE2, and astrocytic and neuronal markers (AQP4 and NeuN) remained unchanged between light and dark phases (Fig. S10F-I).

Together, these findings identify circadian phase as a major source of transcriptional heterogeneity in brain pericytes and support the existence of distinct light- and dark-associated molecular states.

## Discussion

The brain capillary network supports diverse physiological functions across distinct anatomical and functional contexts. Pericytes, which regulate BBB integrity, vascular stability, basal vascular tone, and cerebral blood flow^1,2,4,37^ are therefore likely to require context-dependent molecular organization. Our findings demonstrate that adult mouse brain pericytes exhibit spatially organized and dynamically regulated molecular heterogeneity associated with vascular topology, anatomical region, aging, ischemia, and circadian phase. These findings suggest that pericyte molecular phenotype cannot be understood independently of vascular position and physiological context.

Reanalysis of three published adult mouse brain vascular transcriptomic datasets supported the presence of molecular heterogeneity among brain pericytes. However, the pericyte populations identified across datasets were not identical, suggesting that the analyzed transcriptomic datasets do not yet provide sufficient coverage to define a stable, universal brain-wide taxonomy of pericyte populations. Rather than supporting a small number of universally conserved pericyte subtypes, our analyses point to recurrent but incompletely resolved molecular diversity. This is clearly seen in the hypothalamic dataset from Smith et al.^19^, in which focused sampling of approximately 3,000 pericytes revealed multiple transcriptionally distinct pericyte populations. By contrast, in the multi-region Yao et al. dataset^31^, pericytes from different anatomical regions did not segregate into region-specific clusters, and the number of pericytes per region was relatively low (e.g., 152 hypothalamic pericytes, S. Table 4), limiting assessment of local pericyte diversity at comparable depth (Fig.3). These observations suggest that adult brain pericytes remain under-sampled in current single-cell RNA-seq datasets and that the full profile of pericyte heterogeneity is likely richer than presently appreciated.

The discrepancy between transcriptomic and tissue-based analyses highlights the importance of spatial validation. In the Vanlandewijck et al. dataset ^17^, approximately half of the pericytes were assigned to an *Ace2*-enriched cluster. In contrast, protein-level mapping showed that ACE2*^high^* pericytes were more frequent across most brain regions. This difference may reflect differences in the anatomical regions and vascular territories represented in the dissociated cell suspension, or differential recovery of molecularly distinct pericytes during tissue dissociation. Thus, transcriptomic analyses alone may not have faithfully captured the anatomical abundance and spatial organization of pericyte phenotypes in intact brain vasculature.

Our protein- and mRNA-level analyses show that adult brain pericyte heterogeneity is detectable in situ. ACE2 expression varied among capillary pericytes despite relatively uniform expression of canonical pericyte markers such as CD13 and PDGFRB, allowing pericytes to be stratified into ACE2*^high^* and ACE2*^low^* groups for spatial analysis. In addition, *Igf2*, *Spp1*/OPN, and CASQ2 showed heterogeneous expression among pericytes in brain tissue. The heterogeneous expression of *Igf2* is particularly interesting in light of recent evidence implicating pericyte-derived IGF2 in hippocampal memory formation^36^, suggesting that molecularly distinct pericytes may contribute differently to local neurovascular or parenchymal signaling. Among the identified markers, CASQ2 was particularly notable because both its expression and subcellular organization varied across physiological conditions, indicating that pericyte heterogeneity extends beyond transcriptional differences to dynamic alterations at the protein level.

Our analyses focused on individual markers, but pericyte heterogeneity is likely combinatorial, with phenotypes defined by combinations of molecular features rather than by single marker genes. A comprehensive protein-level classification of brain pericytes will likely require multiplexed imaging approaches that simultaneously quantify multiple markers within intact vascular networks. Such approaches may reveal additional pericyte phenotypes and clarify how marker combinations relate to vascular topology, physiological state, and disease.

Beyond molecular diversity and regional variation, our analyses revealed that pericyte heterogeneity is spatially organized within the cortical vascular network. Although edge-based quantification does not assign ACE2 expression to individual pericyte somata, it enables large-scale assessment of ACE2-associated positional bias while avoiding ambiguous assignment of branch-point pericytes to single vessel segments. By combining large-volume vascular reconstruction with edge-based ACE2 quantification, this analysis provides a spatially resolved view of pericyte-associated molecular heterogeneity that is not captured by cell-type classification alone. ACE2*^high^* vessel edges were enriched near ASMA-positive arterioles, whereas lower ACE2 expression was more frequent in intermediate and more distal capillary segments (Fig. 2D, F and Fig. S2H), indicating that ACE2-associated molecular heterogeneity is not randomly distributed across the cortical microvascular network. Consistent with this concept, Wang et al. recently provided spatial evidence that molecularly defined human pericyte states associate with distinct vascular contexts^25^. Here, three-dimensional vascular reconstruction further relates pericyte-associated molecular heterogeneity to topological position within intact cortical microvascular networks. Topological position may represent an important dimension of pericyte heterogeneity, as increasing evidence indicates that pericytes occupying different locations along the vascular tree differ in their physiological properties and responses to injury ^5,38–40^. The observed positional bias suggests that ACE2-associated heterogeneity may reflect local vascular context, including differences in network architecture, flow-related forces, or metabolic demand, and could potentially influence local angiotensin peptide processing; however, these parameters were not examined. No relationship was detected between capillary diameter and ACE2 expression (Fig. S2E-G), indicating that local differences in capillary caliber did not simply explain ACE2-associated positional bias. Because these measurements were performed in fixed tissue, they do not address whether ACE2 expression is related to dynamic changes in vascular tone in vivo. The spatial organization of molecularly distinct pericytes raises the possibility that they may contribute differently to capillary function, including blood flow regulation and BBB maintenance, a hypothesis that warrants further investigation. Future application of this analytical framework to multiple animals and experimental conditions will enable assessment of whether these spatial patterns are preserved or remodeled during aging and disease.

The relative abundance of ACE2*^high^* and ACE2*^low^* pericytes changed during aging (Fig. 5 A-C). Furthermore, CASQ2 organization was altered upon aging (Fig. 5D-E), following acute ischemia (Fig. 6) and across circadian phase (Fig. 7E-H), with circadian phase emerging as a major contributor to transcriptional heterogeneity among pericytes. These findings extend previous observations that brain pericytes undergo developmental and injury-associated changes^41,42^ and further indicate that pericyte molecular phenotypes can also vary with aging and circadian phase. Loss of pericytes has been described during aging and in neurodegeneration.^12^ However, we did not detect reduced cortical pericyte numbers in 14–17-month-old mice, consistent with Berthiaume et al., who reported no major changes in the structure of the upper cortical capillary network or pericyte soma abundance in 18–24-month-old mice^43^. Thus, the increased proportion of ACE2*^low^* pericytes in the aged cortex is more consistent with phenotypic remodeling than with the selective loss of ACE2*^high^* pericytes. Interestingly, the aged cortex also showed a broader laminar distribution of ACE2*^low^* pericytes (Fig. 5B), indicating that molecular heterogeneity within the cortical pericyte population becomes less spatially segregated during aging. Whether molecularly distinct pericyte populations, including ACE2-stratified groups, differ functionally or exhibit differential vulnerability during aging remains unresolved. This is particularly relevant in light of human studies describing selectively vulnerable pericyte populations in Alzheimer’s disease.^14^ Interestingly, differential *ACE2* expression in human pericyte subtypes can be observed (Fig. S11)^14^, although whether this corresponds to differential ACE2 protein expression remains unknown. Together, these observations indicate that at least some aspects of pericyte heterogeneity reflect dynamic physiological states that can be remodeled over timescales ranging from hours to months. Among the markers examined, CASQ2 showed the greatest sensitivity to physiological contexts, prompting further consideration of its potential functional significance.

CASQ2 is a calcium-buffering protein^44^, and it may represent a molecular feature of pericytes that is particularly sensitive to physiological state, as its expression and subcellular organization varied across circadian phase, aging, and acute ischemia. Although we did not directly measure calcium handling, these findings suggest that CASQ2-positive pericytes may differ in their capacity to buffer intracellular calcium fluctuations under changing physiological or pathological conditions. Recent in vivo calcium imaging studies further demonstrate that pericyte calcium activity is dynamically regulated across vigilance states and differs between brain regions, supporting the concept that calcium-handling mechanisms are physiologically regulated in adult brain pericytes.^45^ Given the established role of calcium signaling in pericyte contractility and ischemia-induced capillary constriction^38,39,46^, CASQ2 immunoreactivity may reflect differential calcium handling in pericytes. Such changes could be adaptive, helping pericytes buffer acute calcium fluctuations, or maladaptive, contributing to altered capillary tone after injury.

The enrichment of *Casq2* and stronger CASQ2 immunostaining in light-phase pericytes are also notable because *Casq2* has been identified as a circadian-regulated gene in cardiomyocytes^47^, where CASQ2 contributes to intracellular calcium handling under catecholamine/cAMP-associated stress^48^. Although the role of CASQ2 in brain pericytes remains unknown, these observations raise the possibility that circadian regulation of calcium storage mechanisms contributes to temporal variation in pericyte physiology. Future studies combining pericyte-specific calcium imaging, vascular function measurements, and CASQ2 perturbation will be required to determine how CASQ2 contributes functionally to pericyte responses during physiological activity, aging, or ischemic injury.

The circadian regulation of the neurovascular unit is increasingly recognized^49^. Daily oscillations in BBB function and vascular homeostasis have been reported^50^; previous studies have implicated BMAL1-dependent pathways in pericyte dysfunction and BBB integrity^51^, and in vitro work suggests that intrinsic pericyte clocks can influence endothelial circadian synchronization and angiogenic responses^52^. Our findings extend these observations by showing that circadian phase represents a major axis of transcriptional heterogeneity in adult brain pericytes in vivo. Importantly, the light–dark separation was not limited to core clock genes but extended to broader functional modules. Dark-phase pericytes were enriched for canonical clock-output genes, as well as genes linked to metabolism, lipid transport, extracellular matrix proteostasis, and stress-response pathways. In contrast, light-phase pericytes showed higher expression of genes associated with calcium-store regulation, ion handling, membrane transport, translation, and extracellular or secretory regulation. These findings suggest that brain pericytes occupy distinct light- and dark-associated transcriptional states, potentially reflecting time-of-day-dependent requirements for metabolic support, vascular stability, barrier regulation, and calcium homeostasis. Whether circadian state transitions are a general feature of pericytes across organs or are particularly pronounced in the neurovascular unit remains an important question for future studies. Notably, we did not detect a comparably strong circadian transcriptional signature in endothelial cells (Fig. 7B). This is consistent with previous work showing that although endothelial BMAL1 regulates circadian BBB function, relatively few endothelial transcripts exhibit robust circadian oscillations^53^.

Collectively, our data demonstrate that molecular heterogeneity is a fundamental feature of adult brain pericytes. Our findings suggest that this heterogeneity comprises both relatively stable spatial phenotypes associated with vascular architecture and dynamically regulated molecular states that change with physiological context. Thus, pericytes exhibit molecular phenotypes that vary by anatomical region, vascular network position, and physiological state. Our findings offer a spatial and dynamic perspective on pericyte heterogeneity that may help understand how the brain capillary network serves diverse functional demands.

## Materials and Methods

### Mice

The following strains and genetically modified mice were used: C57BL/6J, B6 - *Atp13a5^em(2A-CreERT2-IRES-tdTomato)^* (in short, *Atp13a5*-tdTomato)^54^. 2-month-old to 17-month-old mice of both sexes were used. The stroke surgery was performed on 2-3-month-old C57BL/6J mice. Mice were housed in type 2L cages with individual ventilation under specific-pathogen-free conditions and a 12-h light/dark cycle. The cage environment included a red polycarbonate house and shredded paper nesting material. To collect tissue from the light and dark phases, animals were deeply anesthetized and perfused at Zeitgeber Time (ZT) 4 and ZT16. Experimental procedures were approved by the Cantonal Veterinary Office Zurich (ZH039/2024, ZH194/2020, ZH165/2019) or the Institutional Animal Care and Use Committee at Seattle Children’s Research Institute (Protocol IACUC00419).

### Single-cell RNA-seq data analysis

We obtained gene expression profiles of mouse brain vascular cells from three scRNA-seq datasets: Vanlandewijck et al.^17^, Yao et al.^31^, and Smith et al.^19^ (S.Table 1). We chose these datasets because of their purity and, in particular, the large number of vascular cells. In addition, the Yao et al dataset^31^ contains information on the anatomical regions. All mouse brain scRNA-seq datasets were analyzed using Seurat (v. 4.0.3)^55–58^. In the Vanlandewijck et al^17^ dataset, we identified three endothelial cell (EC) clusters (arterial, venous and capillary), pericytes, smooth muscle cells (SMC), PVMs, PVFs and astrocytes (S.Table 2). In total, these endothelial and mural cell clusters from Vanlandewijck et al. contain 2928 cells and were used for further analysis. The Yao et al. dataset (PMID: 37034735) also contains non-vascular cells, such as neurons and oligodendrocytes, from multiple anatomical regions. We extracted only the clusters expressing endothelial cell and mural cell markers, including *Anpep*, *Rgs5*, *Pdgfrb*, *Atp13a5*, *Pecam1, Cldn5*, *Mfsd2a*, *Slc16a1*, and *Tfrc*. In total, these endothelial and mural cell clusters in Yao et al. contain 79’040 cells. After removing data points expressing neuronal marker (*Rbfox3*), microglia marker (*Tmem119*), oligodendrocyte marker (*Mbp*), and astrocyte marker (*Aqp4*). 19’631 cells, including 4’183 mural cells and 7’548 endothelial cells, remained for further analysis (S.Table 1). Detailed cell counts utilized for downstream analyses, broken down by cluster, brain region, sex, and light/dark phase, are provided in Supplementary Table 4. The Smith et al. dataset^19^ contained EYFP-expressing FACS-sorted cells from the hypothalamus of *Gipr*Cre-EYFP and *Glp1r*Cre-EYFP mice. We reanalyzed the *Gipr*Cre-EYFP cells, which are mainly mural cells and a small population of endothelial cells (GSE199301). A total of 3294 cells were used for further analysis (S.Table 1). Notably, while the Smith et al. dataset included cells from both sexes (4 male and 24 female mice; GSE199301), the recovered vascular cells were highly sex-imbalanced, with most captured cells originating from male tissue (Fig. S4A). To avoid clustering artifacts due to sex imbalance and maximize cell yields, subsequent downstream analyses were restricted to the male data. The summary and cell numbers analyzed in this study across the three publicly available datasets are listed in Supplementary Tables 1 - 4.

### The inference of transcription factor (TF) activities and cell-cell communication

Two ligand-receptor pair algorithms, NATMI (Network Analysis Toolkit for Multicellular Interactions)^59^ and CellChat^60^, were applied to compute the probability of cell-cell interactions. Both algorithms calculate the probability of interactions based on the mean expression of ligand and receptor genes at the cell cluster level. NATMI and CellChat applied different scoring systems and ligand-receptor interaction databases (connectomeDB2020 (PMID: 29443965) and CellChatDB). The expression profile of mouse brain vascular cells was obtained from Vanlandewijck et al.^17^ We created the input data for the NATMI and CellChat algorithms based on the cell composition of the capillary. Only pericyte subtypes, astrocytes, and capillary EC profiles were used for interactome analyses. The input data, including gene expression data and cell label metafile, were generated as described in NATMI (https://github.com/asrhou/NATMI) and CellChat (v. 1.1.3)^60^. For ligand-receptor interaction analysis, NATMI was executed using the default ligand-receptor connection database (--interDB lrc2p) and configured for mouse data (--species mouse). The analysis was performed in Python (v. 3.8.7) utilizing pandas (v. 1.0.3), XlsxWriter (v. 1.3.7), xlrd (v. 1.2.0), seaborn (v. 0.11.1), igraph (v. 0.8.3), NetworkX (v. 2.5), PyGraphviz (v. 1.7), Graphviz (v. 0.16), bokeh (v. 2.2.3), and holoviews (v. 1.14.1). Parallel cell-cell communication modeling was conducted using CellChat. First, overexpressed ligands and receptors were identified for each cell type using a Wilcoxon rank-sum test with a significance threshold of *p<0.05*. CellChat then calculated communication probabilities by evaluating the average expression values of a given ligand in a sender cell population relative to its cognate receptor in receiver cell populations, while accounting for multi-subunit cofactors. This analysis was implemented in R (v. 4.1.0) using the packages CellChat, NMF (v. 0.23.0), circlize (v. 0.4.14), ComplexHeatmap (v. 2.11.1), and umap (v. 0.2.8.0). DecoupleR (v 2.9.7)^61^ was applied to infer transcription factor (TF) activity.

### Immunohistochemistry

Deeply anesthetized mice were perfused with phosphate-buffered saline (PBS) for 1–2 min, followed by perfusion with 4% paraformaldehyde (PFA) in PBS (pH 7.2) for 5 min. Brains were collected and post-fixed in 4% PFA in PBS (pH 7.2) at 4 °C for 6 hours. Sixty-μm-thick tissue sections were cut using a vibratome (Leica VT1000S). Floating brain sections were incubated in the blocking/permeabilization solution (1% bovine serum albumin (BSA), 0.5% Triton X-100 in PBS) at 4 °C, followed by incubation in primary antibody solution for 3 days at 4 °C, and subsequently in secondary antibody solution overnight at 4 °C. Following primary antibodies and dilutions were used: anti-CD13 (Biorad, MAC2183, 1:100), anti-collagen IV (Biorad, 2150-1470, 1:600), anti-ASMA-FITC (Sigma-Aldrich, F3777, 1:100), anti-ACE2 (R&D systems, AF3437, 1:100), anti-PDGFRB (eBioscience, 14-1402), anti-osteopontin (R&D systems, AF808, 1:100), anti-CASQ2 (Invitrogen, PA1-913, 1:100), anti-GFAP (Novus Bio, NBP1-05198, 1:600), anti-AQP4 (Novus Bio, NBP1-87679, 1:100), anti-NeuN (Novus Bio, NBP3-05554, 1:100). Secondary antibodies conjugated to fluorophores suitable for multiple labelling were obtained from Jackson ImmunoResearch and used at dilution 1:600. Tissue sections were incubated with DAPI (4’,6-Diamidino-2-phenylindole dihydrochloride) solution (Sigma-Aldrich, D9542, stock 10mg/ml, 1:10000) to label nuclei for 7 minutes at RT, washed and mounted in ProLong Gold or ProLong Diamond Antifade mounting medium (Life Technologies, P36930 and P36965).

### RNA hybridization combined with immunohistochemistry

For simultaneous spatial detection of transcripts and protein targets within the intact neurovascular unit, an optimized immunofluorescence (IF) and multi-signal amplification RNA in situ hybridization (ISH) protocol was performed using the rnaMUSE platform (arcoris bio). To preserve RNA integrity, all buffers were prepared using RNase-free water and supplemented with recombinant RNase inhibitors (New England Biolabs, M0314S). Following deep anesthesia, mice were transcardially perfused with ice-cold PBS, then with 4% PFA in PBS (pH 7.2). Collected brains and hearts were post-fixed overnight in 4% PFA in PBS (pH 7.2) at 4 °C with gentle agitation. Sixty mm tissue sections were cut using a vibratome (Leica VT1000S). Free-floating sections were incubated overnight at 4 °C in freshly prepared MUSE blocking buffer (MUSE-BB: 1% RNase-free BSA, 2% Tween-20 in PBS). Sections were incubated for 2 days at 4 °C on a shaker with primary antibodies diluted in MUSE antibody buffer (MUSE-BB: 0.5% RNase-free BSA, 1% Tween-20 in PBS). Sections were washed with MUSE antibody buffer and post-fixed in 4% PFA at RT to stabilize primary antibody binding before hybridization. Sections were washed with PBS, equilibrated in 2X SSC buffer containing 0.1% Tween-20, and pre-hybridized in pre-warmed hybridization buffer containing RNase inhibitors for 2 h at 37 °C in a humidified chamber. Custom rnaMUSE oligonucleotide probes targeting specific transcripts (*Casq2, Igf2, Spp1, Ace2*) were diluted 100 times in fresh, pre-warmed hybridization buffer. Sections were incubated with the diluted probe overnight at 37 °C with continuous shaking. The unbound probe was removed by washing with pre-warmed wash buffer. All subsequent steps were executed at RT. Next, sections were incubated in enhancement buffer under agitation. The first amplification step was initiated by adding a 1:1 mixture of Muse A and Muse B reagents for 4 h, followed by washes. The second amplification cascade was performed using a 1:1 mixture of Muse C and Muse D reagents for 4 h, and the sections were washed extensively. Sections were blocked in MUSE Blocking Buffer at RT, followed by overnight incubation at 4 °C with fluorophore-conjugated secondary antibodies, and counterstained with DAPI shortly thereafter. Sections were mounted on glass slides using ProLong Diamond Antifade mounting medium (Life Technologies, P36965). Oligonucleotide probes are listed in Supplementary Table 5. Tissue sections, after immunostaining or RNA hybridization combined with immunohistochemistry, were imaged using the slide scanner Zeiss Axio Scan.Z1 (Zeiss) and a confocal microscope (Leica SP8 and Evident Fluoview 4000). Image processing was done using ImageJ2 (v2.16.0), Zeiss ZEN Lite (v. 3.4), and QuPath (v. 0.3.2).

### Ischemic stroke model

The *in situ* thromboembolic stroke model was used to mimic the clinical scenario of stroke according to a published protocol.^62,63^ Briefly, thrombin was injected via a micropipette into the M2 segment of the middle cerebral artery (MCA) of anesthetized mice to induce a clot *in situ*. Simultaneous monitoring of cerebral blood flow using laser speckle imaging (FLPI, Moor Instruments, UK) was used to confirm a reduction in blood flow after blood clot formation. Sham-operated control mice underwent an identical surgical exposure and imaging regimen. Mice were sacrificed 4 h after stroke induction, before spontaneous reperfusion in the hyperacute phase.

### IDISCO tissue clearing and 3D rendering

Deeply anesthetized 1-month-old *Atp13a5*Cre-tdTomato mice were perfused with PBS for 1–2 min, followed by perfusion with 4% PFA in PBS (pH 7.2) for 5 min. Brains were collected and post-fixed in 4% PFA in PBS (pH 7.2) at 4 °C for 6 hours. The immunolabeling was performed following the iDISCO protocol.^64^ The brain tissue sections were incubated with antibodies anti-ASMA-FITC (Sigma-Aldrich, F3777, 1:200), anti-ACE2 (R&D systems, AF3437, 1:200) and anti-collagen IV (Biorad, 2150-1470, 1:500) for one week at 37 °C with rotating and then further incubated with the secondary antibodies conjugated to fluorophores suitable for multiple labelling obtained from Jackson ImmunoResearch (1:1000) for one week at 37 °C with rotating. Brain tissues were cleared following the FDISCO protocol^65^ to reduce peroxide-induced quenching. Dehydration was performed using a graded series of tetrahydrofuran solutions adjusted to pH 9.0: 50% (v/v), 70% (v/v), 80% (v/v), and 100% (v/v) (three times each). Each incubation step lasted at least 12 hours and was carried out at 4°C with continuous vertical rotation using a tube rotator. After FDISCO, the tissues were stored in dibenzyl ether (DBE) at 4 °C. Cleared tissues were placed in a confocal dish (VWR, 75856-742) filled with DBE and imaged using an in-house-built benchtop^66^ mesoscale selective plane illumination microscope with a Mitutoyo M Plan Apo 5x objective/NA=0.14.

### Image analysis

#### Measurements of the vessel diameter and ACE2, CD13 intensity

Quantification of vessel diameter and protein intensity was performed on images acquired with a Leica SP8 confocal laser scanning microscope (Leica Microsystems GmbH, 20X objective/NA=0.75, 63X objective/NA=1.4). Images were acquired from the deep brain, cortex, and hippocampus. Vessel diameter was determined using VasoMetrics, a FIJI macro based on collagen IV staining.^67^ The regions of interest (ROIs) were manually annotated on pericyte somata based on CD13 intensity, in parallel with vessel diameter measurements. For each image, 10-15 ROIs (i.e., pericytes) were selected, with a total of 417 ROIs obtained from three mice. In addition, the corresponding ACE2 and CD13 intensities were measured in FIJI using the default measurement function.

#### Pericyte quantification

Images were acquired using an AxioScan.Z1, and the cell detection function in QuPath (v. 0.3.2) was used to identify pericytes in the raw images (.czi files). First, pericytes were identified by CD13 or tdTomato expression. The following parameters were applied to detect pericytes: requested pixel size - 0.5, background radius - 10, median filter radius - 0, Sigma - 3, minimum area - 50, and maximum area - 1000. CD13 signal intensity greater than 2000 or tdTomato signal intensity greater than 2500 was used to identify a pericyte soma. Second, ACE2 intensity was used as a parameter to distinguish pericyte groups. Two groups (ACE2*^high^* and ACE2*^low^*) were separated based on a threshold of ACE2 intensity, defined as one-third of the entire ACE2 intensity distribution, derived from the overview histogram of intensity values and cell counts calculated by QuPath. Cells expressing ACE2 higher than this threshold were defined as ACE2*^high,^* and those lower than this threshold were defined as ACE2*^low^* pericytes. Brain sections processed and immunostained simultaneously were analyzed using the same ACE2 threshold levels. The percentages of ACE2*^high^* and ACE2*^low^* pericytes were identified across different brain regions, manually annotated based on the Allen Brain Atlas (mouse.brain-map.org), including cortical regions (anterior cingulate area, somatomotor area, somatosensory area, subplate), hypothalamus, thalamus, septal area, corpus callosum, and hippocampus.

#### Annotation and segmentation

We analyzed ACE2-positive edge distribution in the cortical region of a 3-month-old brain hemisphere. The vascular tree was segmented from the tdTomato channel using a fine-tuned vesselFM model^68^. To identify arteries within this segmentation, we trained a 3D nnU-Net (ResEnc-L, 3d_fullres)^69^ from scratch on the tdTomato segmentation and raw ASMA channel as inputs, using 10 expert-annotated cutouts (5-fold cross-validation, mean Dice 0.67), then applied it to the full light-sheet volume. Voreen’s vessel graph extraction algorithm^70^ was used to extract a skeleton graph (nodes: vessel junctions; edges: individual capillary or vessel segments). Each segmentation voxel was assigned to its nearest skeleton edge by a topology-respecting, multi-source watershed seeded at every skeleton voxel, where edges with at least 20% arterial voxel fraction were labeled as artery. To classify ACE2 positivity, we trained an L2-regularized logistic regression classifier on each edge’s background-corrected ACE2 intensity (raw intensity minus a locally estimated background) using 800 expert-labeled capillaries (5-fold cross-validated AUC 0.84), then classified all edges as ACE2-*high* or -*low*. The population was restricted to cortical capillaries via an expert-drawn mask. Cumulative path distance from each cortical capillary to the nearest artery segment was computed by a multi-source Dijkstra algorithm along the skeleton graph. To test whether ACE2*^high^* edges are enriched near arteries relative to the mid-capillary bed, we divided the artery-to-capillary distance range into [3/5] equal-width bands. We computed a 2×2 contingency table (near band [bin 1] vs. middle band [center bin] by ACE2 status) within each artery tree. Tree-level log-odds ratios were pooled by DerSimonian–Laird random-effects meta-analysis, accounting for between-territory heterogeneity (I²) rather than assuming a single shared effect.

#### Quantification of CASQ2 positive area

Quantification of CASQ2 positive area was performed on images acquired with a Leica SP8 confocal laser scanning microscope (Leica Microsystems GmbH, 20X objective/NA=0.75A). At least three images per region were acquired from young (*n = 5*) and aged animals (*n = 5*), ischemic and contralateral cortices of stroke mice (*n = 4*), and the corresponding ipsilateral cortex of sham-operated controls (*n = 4*). Confocal Z-stacks spanning a total optical thickness of 18 mm were used to generate maximum-intensity projection images in Fiji for subsequent analysis. Regions of interest (ROIs) corresponding to the pericyte cell bodies were automatically segmented based on CD13 signal intensity using QuPath (v. 0.3.2). The following parameters were applied to detect pericytes: requested pixel size = 0.5, background radius = 10, median filter radius = 0, Sigma = 3, minimum area = 50, and maximum area = 600. To ensure data integrity, only intact cell bodies that were completely captured within the imaging plane were retained for further analysis. For each image, 15–20 ROIs (pericytes) were selected, resulting in 1190 ROIs obtained across eight stroke-operated mice and 638 ROIs across young and aged animals. The pixel classification function was applied to identify CASQ2-positive and CASQ2-negative areas based on CASQ2 intensity. The total area and percentage area of CASQ2 within pericyte cell bodies were measured. In addition, the corresponding ACE2 and CD13 intensities of the cell body were measured using QuPath.

### Statistical analysis

Statistical analysis was performed using GraphPad Prism (version 11.0; GraphPad Software, La Jolla, CA, USA), R (version 4.6.0 (2026-04-24)), and RStudio (version 2026.05.0+218). Statistical significance between groups was calculated using an unpaired *t*-test with Welch’s correction and one- or two-way ANOVA with Tukey’s multiple comparisons test. Pearson’s chi-squared test was performed to assess the association between pericyte subcluster and tissue collection time (dark and light), with the chi-squared statistic () calculated to determine the effect size. Statistical tests used and the group size (n) are stated in the figure legends. All data are presented as mean ± SD (standard deviation).

## Acknowledgments

The authors acknowledge support from the Center for Microscopy and Image Analysis and the Science IT team (www.s3it.uzh.ch) at the University of Zurich. We thank Liam Sullivan for technical support, Dr. Tania Wyss for tutorials on analyzing single-cell RNA-seq data, Dr. Liqun He for providing the count matrix Excel file for the scRNA-seq data used for NATMI, and Dr. Zhen Zhao for sharing the *Atp13a5*-tdTomato line.

## Author’s contribution

S-F.H. and A.K. conceptualized and designed experiments. S-F.H, S.S., U.M., C.G., M.E.A., H.P., and J.D. performed experiments. L.G. analyzed light-sheet imaging data. B.W., S.W., A.Y.S., and D.R. contributed reagents/resources. A.Y.S. contributed to data interpretation and discussion. S-F.H. and A.K wrote the manuscript. A.K. supervised the study. All authors commented on the manuscript.

