## Supplementary material for "Brain pericytes exhibit spatially organized and dynamically regulated molecular heterogeneity": Figure S1-S10

### Supplementary Figures and Tables

Figure S1. ACE2 expression in brain and retinal mural cells

Figure S2. Spatial distribution and vascular-topological organization of ACE2-defined pericyte groups

Figure S3. Transcriptomic profiles and cell-type-specific marker expression in the Vanlandewijck et al. dataset[17]

Figure S4. Transcriptomic profiles and cell-type-specific marker expression in the Smith et al. dataset[19]

Figure S5. Transcriptomic profiles and cell-type-specific marker expression in the Yao et al. dataset[31]

Figure S6. Inferred transcription factor (TF) activity identifies convergent regulatory programs in brain mural cells from Yao et al. dataset[31]

Figure S7. Spatial validation of molecular heterogeneity in brain pericytes

Figure S8. Age-associated changes in the abundance of ACE2$^{high}$ and ACE2$^{low}$

Figure S9. Acute ischemia does not alter CD13 immunoreactivity or the proportion of ACE2-defined pericyte groups

Figure S10. Transcriptomic analyses and validation of circadian-associated pericyte heterogeneity

Figure S11. Heterogeneous *ACE2* expression in human pericytes.

S. Table 1. Summary of reanalyzed RNAseq datasets

S. Table 2. Vanlandewijck et al. dataset[17]

S. Table 3. Smith et al. dataset[19]

S. Table 4. Yao et al. dataset[31]

S. Table 5. RNA probe sequences

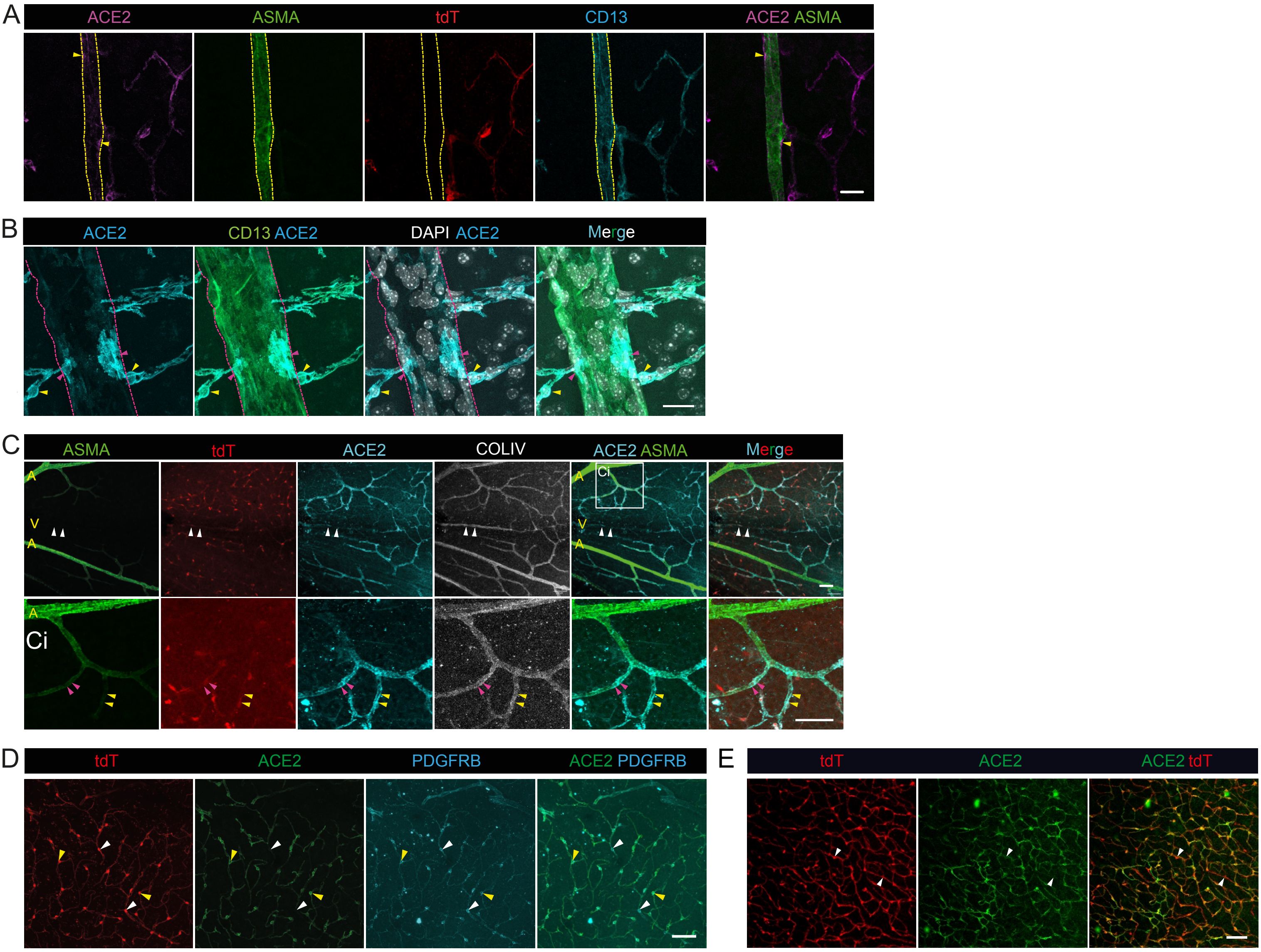
A
ACE2
ASMA
tdT
CD13
ACE2 ASMA
B
ACE2
CD13 ACE2
DAPI ACE2
Merge
C
ASMA
tdT
ACE2
COLIV
ACE2 ASMA
Merge
A
V
Ci
D
tdT
ACE2
PDGFRB
ACE2 PDGFRB
E
tdT
ACE2
ACE2 tdT

**Figure S1. ACE2 expression in brain and retinal mural cells. A** ASMA (in green) positive mural cells (CD13, in cyan) display weak ACE2 (in magenta, yellow arrowheads) immunoreactivity in the cortex of *Atp13a5*-tdTomato mice, where pericytes are genetically labeled (tdT, in red). The dotted yellow line marks the ASMA-positive arteriole. Scale bar, 20 µm. **B** ACE2-positive pericytes (in cyan, yellow arrowheads) extend long, longitudinal processes (pink arrowheads) along a larger venule. Mural cells and pericytes are identified by CD13 immunostaining (in green). The dotted pink line marks the venule. Scale bar, 20 µm. **C** ACE2 (in cyan) immunoreactivity in retinal mural cells in *Atp13a5*-tdTomato mice, where pericytes are genetically labeled (tdT, in red). Upper panels show an overview of the retinal vascular network, with white arrowheads pointing to ACE2-positive mural cells associated with a venule (V). The boxed region (**Ci**) is shown at higher magnification in the lower panels. Pink arrowheads indicate ACE2-positive pre-capillary pericytes that retain ASMA expression, whereas yellow arrowheads indicate adjacent ACE2-positive pericytes in which ASMA expression is strongly reduced, illustrating the transition from ASMA-positive to ASMA-negative mural cells. Collagen IV (in white) labels the vascular basement membrane. A, arterioles. Scale bars, 50 µm. **D** ACE2$^{low}$ pericytes (white arrowheads) and ACE2$^{high}$ pericytes (yellow arrowheads) are positive for the pericyte marker PDGFRB (in cyan) in the cerebral cortex of *Atp13a5*-tdTomato (tdT, in red) mice. Scale bar, 50 µm. **E** The distribution of ACE2$^{low}$ pericytes (white arrowheads) in the *Atp13a5*-tdTomato mouse retinal capillaries. Scale bar, 50 µm. Confocal images are presented as maximum intensity projections.

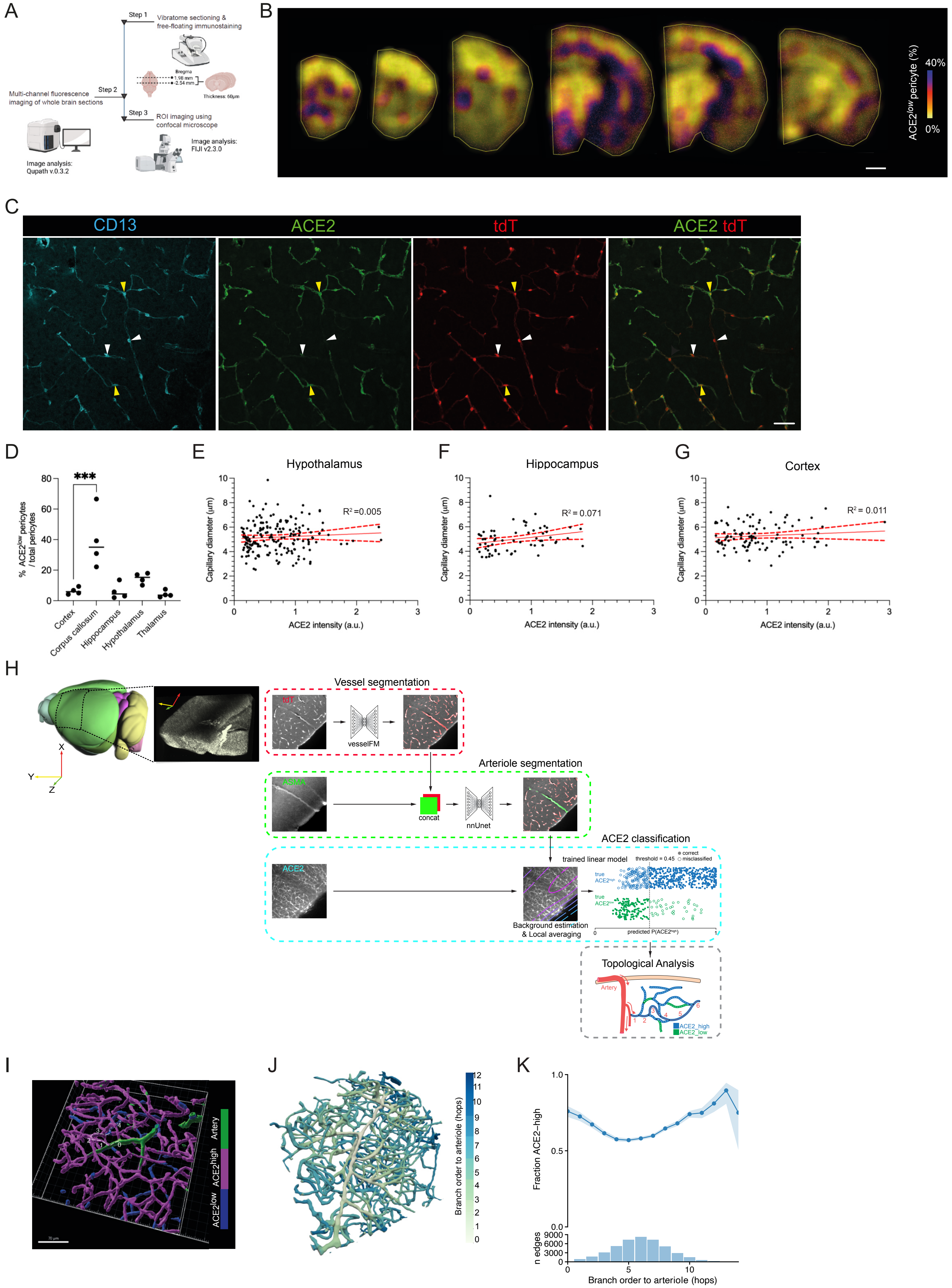
A
Step 1
Vibratome sectioning &
free-floating immunostaining
Bregma
-1.98 mm
-2.54 mm
Thickness: 60μm
Step 2
Multi-channel fluorescence
imaging of whole brain sections
Image analysis:
Qupath v.0.3.2
Step 3
ROI imaging using
confocal microscope
Image analysis:
FIJI v2.3.0
B
ACE2low pericyte (%)
40%
0%
C
CD13
ACE2
tdT
ACE2 tdT
D
***
% ACE2low pericytes / total pericytes
Cortex
Corpus callosum
Hippocampus
Hypothalamus
Thalamus
E
Hypothalamus
R² =0.005
Capillary diameter (μm)
ACE2 intensity (a.u.)
F
Hippocampus
R² = 0.071
G
Cortex
R² = 0.011
H
Vessel segmentation
tdT
vesselFM
Arteriole segmentation
concat
nnUnet
ACE2 classification
ACE2
trained linear model
threshold = 0.45
correct
misclassified
Background estimation
& Local averaging
predicted P(ACE2high)
Topological Analysis
Artery
ACE2_high
ACE2_low
X
Y
Z
I
Artery
ACE2high
ACE2low
70 μm
J
Branch order to arteriole (hops)
K
Fraction ACE2–high
n edges
Branch order to arteriole (hops)

**Figure S2. Spatial distribution and vascular-topological organization of ACE2-defined pericyte groups. A** Experimental workflow for whole-hemisphere protein expression profiling in cortical pericytes. Brain tissue was sectioned coronally (60 µm thick, Bregma 1.98 mm to -2.54 mm) using a vibratome. Free-floating sections underwent multi-channel fluorescence immunohistochemistry for pericyte markers, followed by slide mounting and automated scanning (Zeiss Axio Scan.Z1). Macro-scale spatial profiling was done using QuPath, whereas high-resolution analysis was done using a confocal laser scanning microscope and ImageJ/FIJI. Schematic created with BioRender. **B** Heatmap visualization of ACE2$^{low}$ pericyte proportion (%) across serial coronal brain sections, progressing from anterior to posterior. The color scale ranges from yellow (low proportion of ACE2$^{low}$ pericytes) to blue/purple (high proportion), as indicated by the color bar. Regional heterogeneity in ACE2$^{low}$ pericyte abundance is evident across brain sections, with distinct patterns observed between cortical and subcortical structures. Scale bar, 1 mm. **C** The distribution of ACE2$^{low}$ (white arrowheads) and ACE2$^{high}$ (yellow arrowheads) pericytes in 2–3-month-old *Atp13a5*-tdTomato mice, where pericytes are genetically labeled (tdT, in red). Pan-pericyte marker CD13 is in cyan. Scale bar, 100 µm. **D** The percentage of ACE2$^{low}$ pericytes in different brain regions in *Atp13a5*-tdTomato mice. Each data point represents an individual animal ($n$ = 4 mice). Data are presented as mean ± SD. *** -$p$<0.001 One-way ANOVA with Tukey's multiple comparison test was used to analyze data. **E-F** The vessel diameter does not correlate with the ACE2 expression level in pericytes in the hypothalamus (**E**), hippocampus (**F**), and cortex (**G**). Each data point represents an individual cell; **E**: $n$ = 183 cells from 3 mice, **F:** $n$ = 64 cells from 2 mice, **G**: $n$ = 101 cells from 3 mice. The red line shows the best-fit regression line, and the dashed red lines show the 95% confidence interval. **H** Workflow for the segmentation of volume light-sheet images from an *Atp13a5*-tdTomato mouse cortical hemisphere. 3D schematic of the whole mouse brain showing the cropped cortical volume (black dashed box) and its corresponding high-resolution fluorescence image inset (right). Colored arrows define the spatial coordinate axes (X, red; Y, yellow; Z, green). 3D vessels and arterioles were segmented using tdT and ASMA signals, respectively. The automatic categorization of pericyte subtypes was based on ACE2 expression using a trained linear model. Detailed processing workflows are provided in the Materials and Methods section. **I** 3D volume reconstruction of a vascular tree illustrating the localization of ACE2$^{low}$ pericytes (in blue) and ACE2$^{high}$ pericytes (in purple) from a confocal image. ASMA-positive arterial termini are in green. Scale bar, 70 µm. The white dotted line indicates the branches, and the numbers indicate branch orders.

**J** Branch-order representation of a representative cortical vascular tree, with branch order defined from ASMA-positive arteriolar termini ($0^{th}$ order) and visualized by color intensity (white, proximal; blue, distal). **K** Fraction of ACE2$^{high}$ vessel edges as a function of branch order within the cortical vascular network. The upper plot shows the mean fraction for each branch order, with shading indicating the 95% confidence interval. The lower histogram shows the total number of vessel edges analyzed at each corresponding branch order. ACE2$^{high}$ fractions were higher in low-order branches ($1^{st}$–$3^{rd}$), although the distribution across branch orders was non-monotonic.

Figure S3 Huang et al.

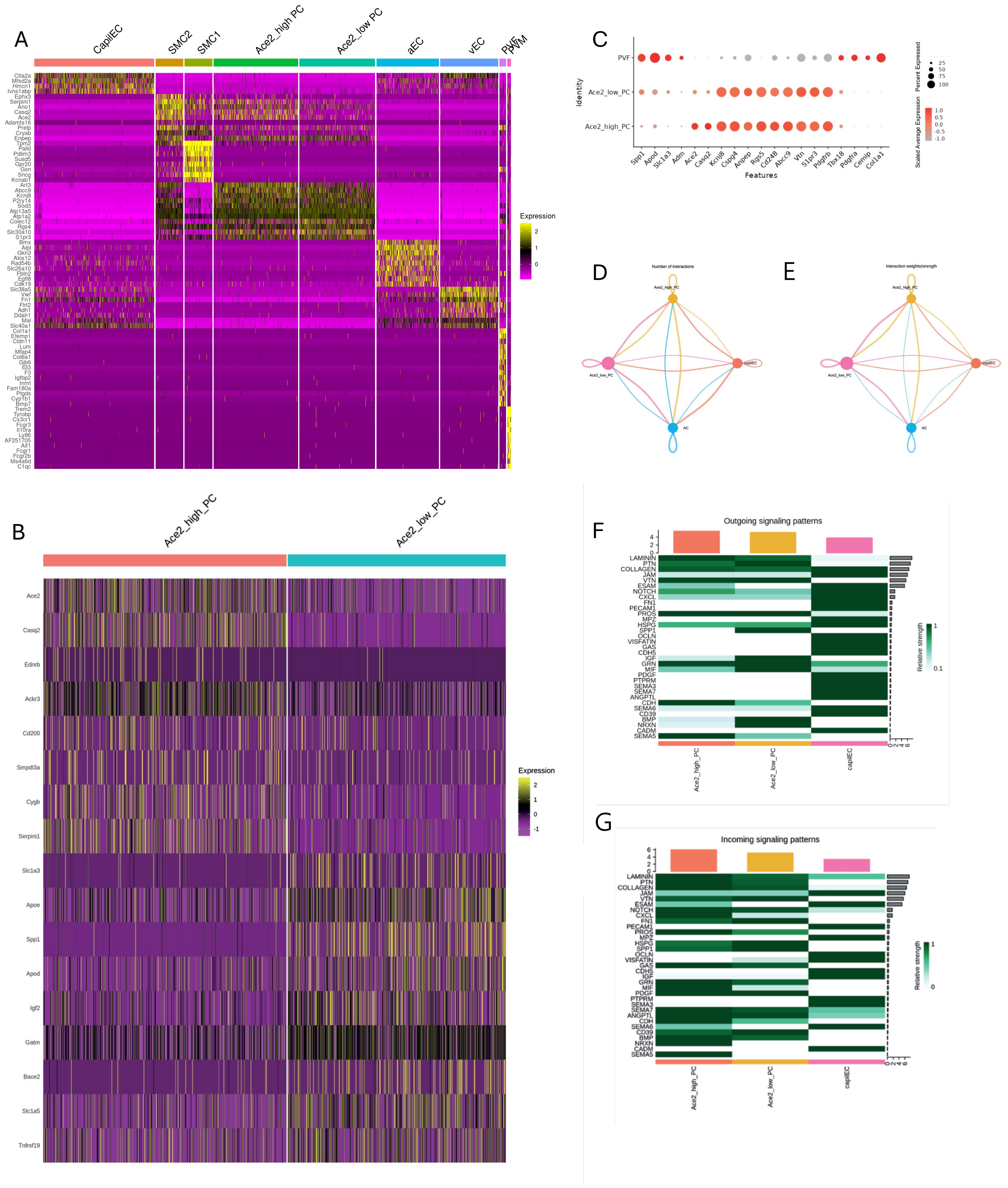

**Figure S3. Transcriptomic profiles and cell-type-specific marker expression in the Vanlandewijck et al. dataset**[17]. **A** Heatmap showing the top 10 differentially expressed genes for each cell cluster. **B** Heatmap displaying the top 10 differentially expressed genes among Ace2_low and Ace2_high pericytes. Color intensity reflects scaled average expression: Magenta and yellow indicate expression of the respective genes (as shown in the legends). **C** Dot plot displaying the scaled average expression of selected marker genes for perivascular fibroblasts (PVF) and pericytes. **D-G** Cell-cell communication analysis using CellChat. Number of significant ligand-receptor pairs between pericyte (PC) subtypes, astrocytes (AC) and capillary endothelial cells (capiEC) (**D**). The edge width is proportional to the indicated number of ligand-receptor pairs. Weighted/strength-directed network between PC subtypes, AC and EC (**E**). Circle sizes are proportional to the number of cells in each cell group and edge width represents the communication probability. Colors indicate specific clusters: Ace2_high pericytes (yellow), Ace2_low pericytes (pink), capiEC (orange) and astrocyte (blue). Heatmap showing the top outgoing (**F**) and incoming (**G**) signaling pathways from Ace2_high and Ace2_low pericytes and capillary endothelial cells (capiEC). The top-colored bar plot shows the total signaling strength of a cell group by summarizing all signaling pathways displayed in the heatmap. The right grey bar plot shows the total signaling strength of a signaling pathway by summarizing all cell groups displayed in the heatmap.

Figure S4 Huang et al.

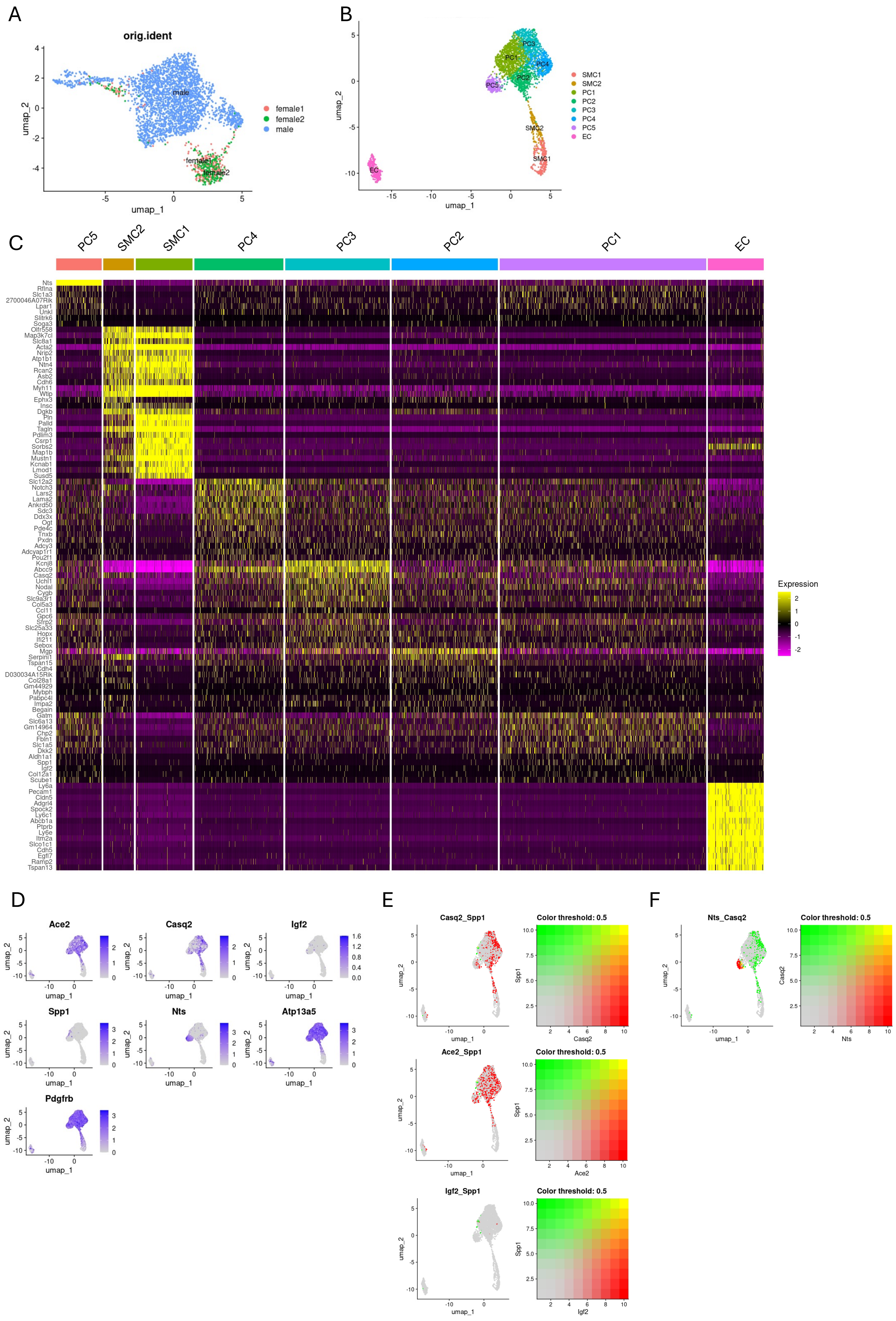

**Figure S4. Transcriptomic profiles and cell-type-specific marker expression in the Smith et al. dataset**[19]. **A** UMAP visualization of the whole Smith et al dataset color-coded by sex, based on annotations provided in the original metadata. **B-D** Transcriptomic profiles and cell-type-specific marker expression in the Smith et al. dataset cells from four males. UMAP visualization at a resolution of 0.6 and colored according to identified cell clusters (**B**). Heatmap showing the top 10 differentially expressed genes for each cluster. Color intensity reflects scaled average expression: Magenta and yellow indicate expression of the respective genes (as shown in the legends) (**C**). Feature plots showing expression of selected genes (*Ace2, Casq2, Igf2, Spp1, Nts*) and pan-pericyte marker genes (*Atp13a5, Pdgfrb*) within the endothelial and mural cell clusters (**D**). **E-F** Bivariate feature plots illustrating the largely non-overlapping expression patterns of selected pericyte marker genes (*Casq2, Spp1, Ace2, Igf2,* and *Nts*) from four males. Each dot represents an individual cell. Green and red indicate expression of the respective genes, whereas yellow denotes cells with strong co-expression of both genes.

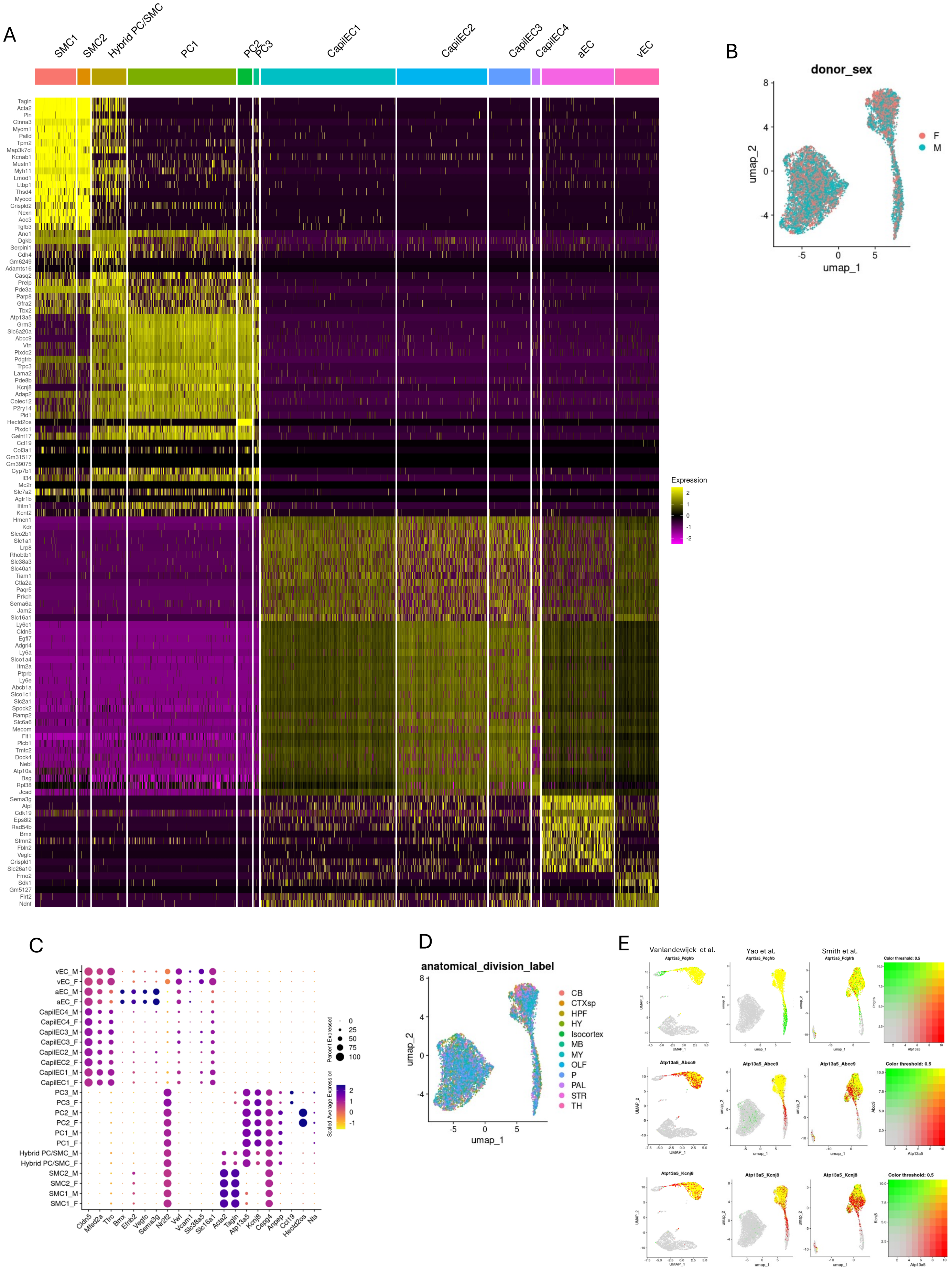
A
SMC1
SMC2
Hybrid PC/SMC
PC1
PC2
PC3
CapilEC1
CapilEC2
CapilEC3
CapilEC4
aEC
vEC
Tagln
Acta2
Pln
Ctnna3
Myom1
Palld
Tpm2
Map3k7cl
Kcnab1
Mustn1
Myh11
Lmod1
Ltbp1
Thsd4
Myocd
Crispld2
Nexn
Aoc3
Tgfb3
Ano1
Dgkb
Serpini1
Cdh4
Gm6249
Adamts16
Casq2
Prelp
Pde3a
Parp8
Gfra2
Tbx2
Atp13a5
Grm3
Slc6a20a
Abcc9
Vtn
Plxdc2
Pdgfrb
Trpc3
Lama2
Pde8b
Kcnj8
Adap2
Colec12
P2ry14
Pld1
Hectd2os
Plxdc1
Galnt17
Ccl19
Col3a1
Gm31517
Gm39075
Cyp7b1
Il34
Mc2r
Slc7a2
Agtr1b
Ifitm1
Kcnt2
Hmcn1
Kdr
Slco2b1
Slc1a1
Lrp8
Rhobtb1
Slc38a3
Slc40a1
Tiam1
Ctla2a
Paqr5
Prkch
Sema6a
Jam2
Slc16a1
Ly6c1
Cldn5
Egfl7
Adgrl4
Ly6a
Slco1a4
Itm2a
Ptprb
Ly6e
Abcb1a
Slco1c1
Slc2a1
Spock2
Ramp2
Slc6a6
Mecom
Flt1
Plcb1
Tmtc2
Dock4
Nebl
Atp10a
Bsg
Rpl38
Jcad
Sema3g
Alpl
Cdk19
Eps8l2
Rad54b
Bmx
Stmn2
Fbln2
Vegfc
Crispld1
Slc26a10
Fmo2
Sdk1
Gm5127
Flrt2
Ndnf
Expression
2
1
0
-1
-2
B
donor_sex
F
M
umap_2
umap_1
C
vEC_M
vEC_F
aEC_M
aEC_F
CapilEC4_M
CapilEC4_F
CapilEC3_M
CapilEC3_F
CapilEC2_M
CapilEC2_F
CapilEC1_M
CapilEC1_F
PC3_M
PC3_F
PC2_M
PC2_F
PC1_M
PC1_F
Hybrid PC/SMC_M
Hybrid PC/SMC_F
SMC2_M
SMC2_F
SMC1_M
SMC1_F
Cldn5
Mfsd2a
Tfrc
Bmx
Efnb2
Vegfc
Sema3g
Nr2f2
Vwf
Vcam1
Slc38a5
Slc16a1
Acta2
Tagln
Atp13a5
Kcnj8
Cspg4
Anpep
Ccl19
Hectd2os
Nts
Percent Expressed
0
25
50
75
100
Scaled Average Expression
2
1
0
-1
D
anatomical_division_label
CB
CTXsp
HPF
HY
Isocortex
MB
MY
OLF
P
PAL
STR
TH
umap_2
umap_1
E
Vanlandewijck et al.
Yao et al.
Smith et al.
Atp13a5_Pdgfrb
Atp13a5_Abcc9
Atp13a5_Kcnj8
Color threshold: 0.5
Pdgfrb
Abcc9
Kcnj8
Atp13a5
UMAP_1
UMAP_2
umap_1
umap_2

**Figure S5. Transcriptomic profiles and cell-type-specific marker expression in the Yao et al. dataset[31]. A** Heatmap showing the top 10 differentially expressed genes for each vascular cluster. Color intensity reflects scaled average expression (Magenta, low; yellow, high). **B** UMAP visualization color-coded by sex based on annotations provided in the original metadata. **C** Dot plot displaying the expression of selected marker genes for endothelial cells (arterial, venous and capillary), subtypes of smooth muscle cells and pericytes, grouped by sex. M, male. F, female. **D** UMAP visualization colored by anatomical regions based on annotations provided in the original metadata. CB, cerebellum; CTX, cerebral cortex; CTXsp, cortical subplate; HPF, hippocampal formation; HY, hypothalamus; IsoCTX, iso cortex; MB, midbrain; OLF, olfactory areas; P, pons; PAL, pallidum; STR, striatum; TH, thalamus. **E** Bivariate feature plots comparing the expression of selected pericyte marker genes. Each dot represents an individual cell. Color intensity reflects gene expression: green and red indicate expression of the respective genes (as shown in the legends), whereas yellow indicates co-expression of both genes.

Figure S6



A

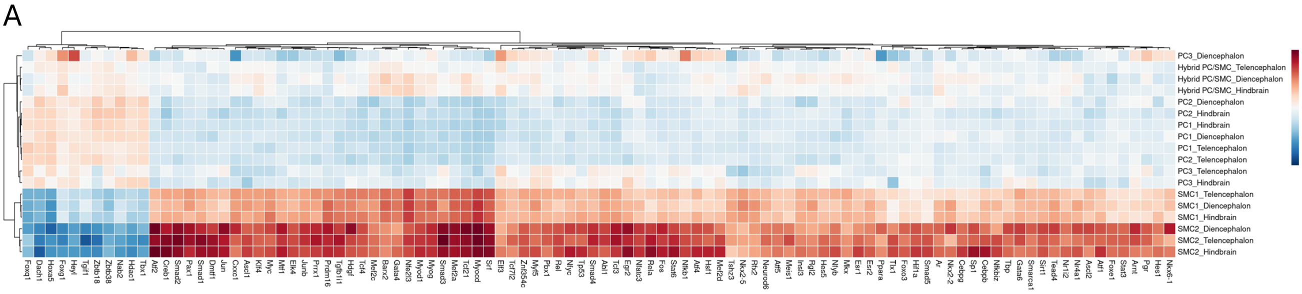
PC3_Diencephalon
Hybrid PC/SMC_Telencephalon
Hybrid PC/SMC_Diencephalon
Hybrid PC/SMC_Hindbrain
PC2_Diencephalon
PC2_Hindbrain
PC1_Hindbrain
PC1_Diencephalon
PC1_Telencephalon
PC2_Telencephalon
PC3_Telencephalon
PC3_Hindbrain
SMC1_Telencephalon
SMC1_Diencephalon
SMC1_Hindbrain
SMC2_Diencephalon
SMC2_Telencephalon
SMC2_Hindbrain
2
1
0
-1
-2

B

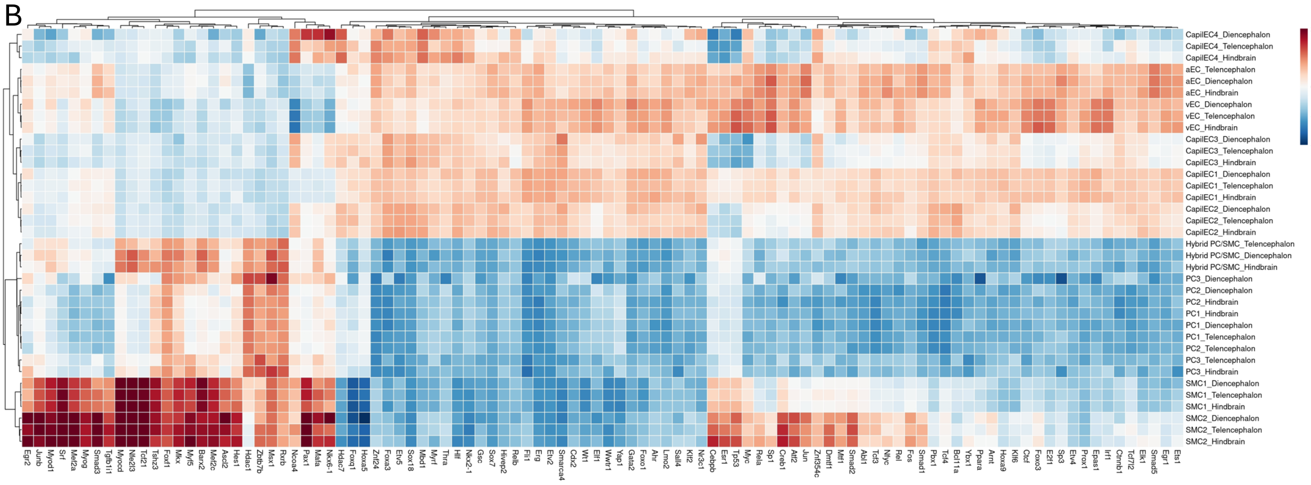
CapilEC4_Diencephalon
CapilEC4_Telencephalon
CapilEC4_Hindbrain
aEC_Telencephalon
aEC_Diencephalon
aEC_Hindbrain
vEC_Diencephalon
vEC_Telencephalon
vEC_Hindbrain
CapilEC3_Diencephalon
CapilEC3_Telencephalon
CapilEC3_Hindbrain
CapilEC1_Diencephalon
CapilEC1_Telencephalon
CapilEC1_Hindbrain
CapilEC2_Diencephalon
CapilEC2_Telencephalon
CapilEC2_Hindbrain
Hybrid PC/SMC_Telencephalon
Hybrid PC/SMC_Diencephalon
Hybrid PC/SMC_Hindbrain
PC3_Diencephalon
PC2_Diencephalon
PC2_Hindbrain
PC1_Hindbrain
PC1_Diencephalon
PC1_Telencephalon
PC2_Telencephalon
PC3_Telencephalon
PC3_Hindbrain
SMC1_Diencephalon
SMC1_Telencephalon
SMC1_Hindbrain
SMC2_Diencephalon
SMC2_Telencephalon
SMC2_Hindbrain
2
1
0
-1
-2

**Figure S6. Inferred transcription factor (TF) activity identifies convergent regulatory programs in brain mural cells from Yao et al. dataset[31]. A** Heatmap of the top 100 differentially active transcription factors in mural cells grouped by anatomical region (telencephalon, diencephalon, and hindbrain). Pericytes and VSMCs cluster primarily by cell type rather than by anatomical region. **B** Heatmap of the top 100 differentially active transcription factors comparing endothelial and mural cell populations across the three anatomical regions (telencephalon, diencephalon, and hindbrain). Pericytes and VSMCs share a common mural cell TF activity signature distinct from that of endothelial cells. Color intensity represents the scaled inferred TF activity (red: high; blue: low).

Figure S7 Huang et al.

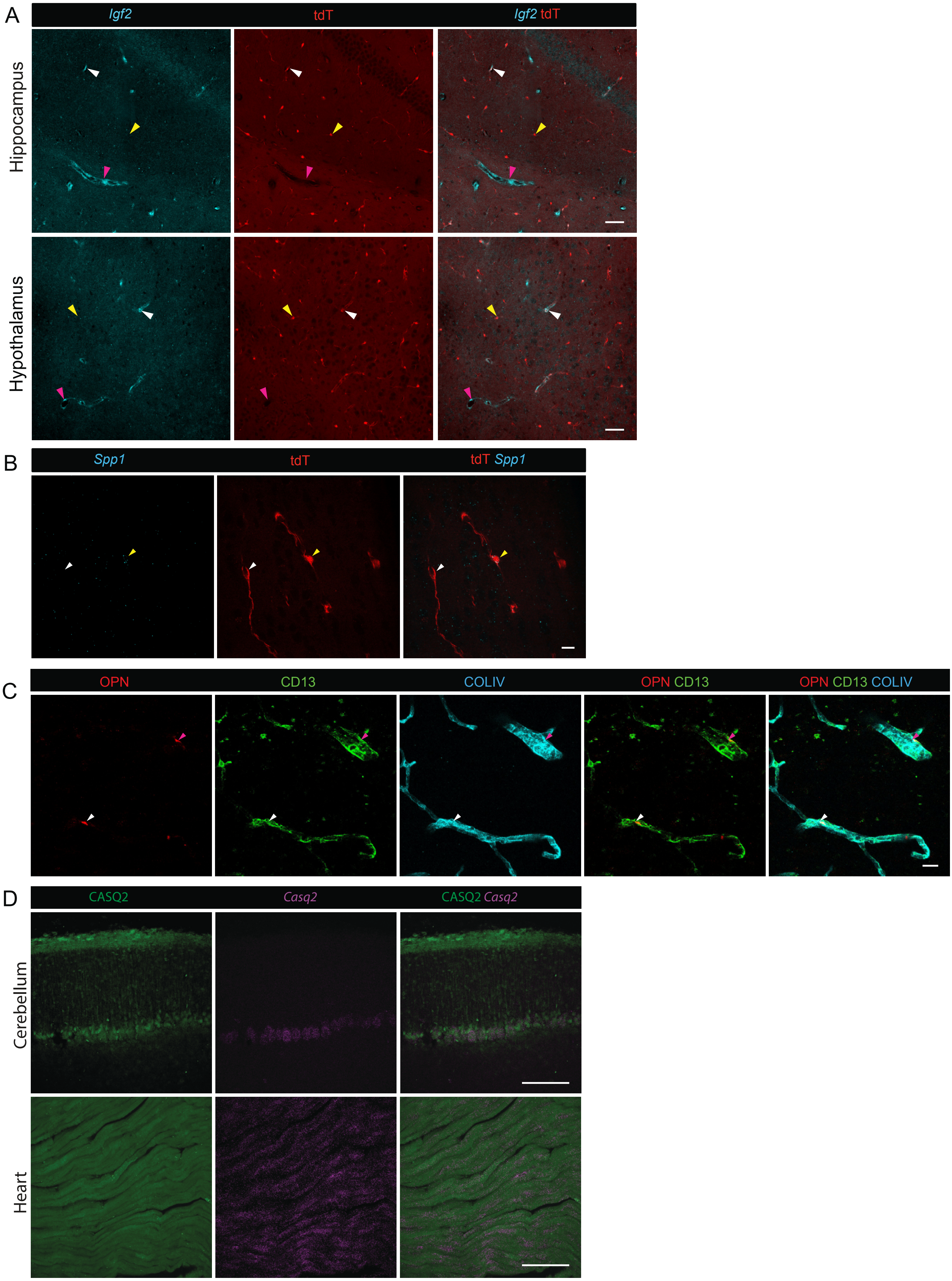

**Figure S7. Spatial validation of molecular heterogeneity in brain pericytes. A** *Igf2* mRNA expression (in cyan) in pericytes (tdT, in red) and perivascular fibroblasts (pink arrowhead) in the hippocampus and hypothalamus of *Atp13a5*-tdTomato mice. White arrowheads indicate *Igf2*-positive pericytes and yellow arrowheads indicate *Igf2*-negative pericytes. Scale bars, 100 µm. **B** *Spp1* mRNA expression (in cyan) in pericytes. A single optical section is presented. White and yellow arrowheads indicate *Spp1*-positive and *Spp1*-negative pericytes, respectively. Scale bar, 10 µm. **C** Osteopontin (OPN; in red) expression in pericytes and perivascular fibroblasts. CD13 (in green) and collagen IV (COLIV, in cyan) are shown for reference. White and pink arrowheads indicate pericytes and perivascular fibroblasts, respectively. Scale bar, 20 µm. **D** Validation of CASQ2 protein (in green) and *Casq2* mRNA (in magenta) expression in wild-type mouse cerebellum and heart. Scale bars, 500 µm. Confocal images in panels A and C–D are presented as maximum intensity projections.

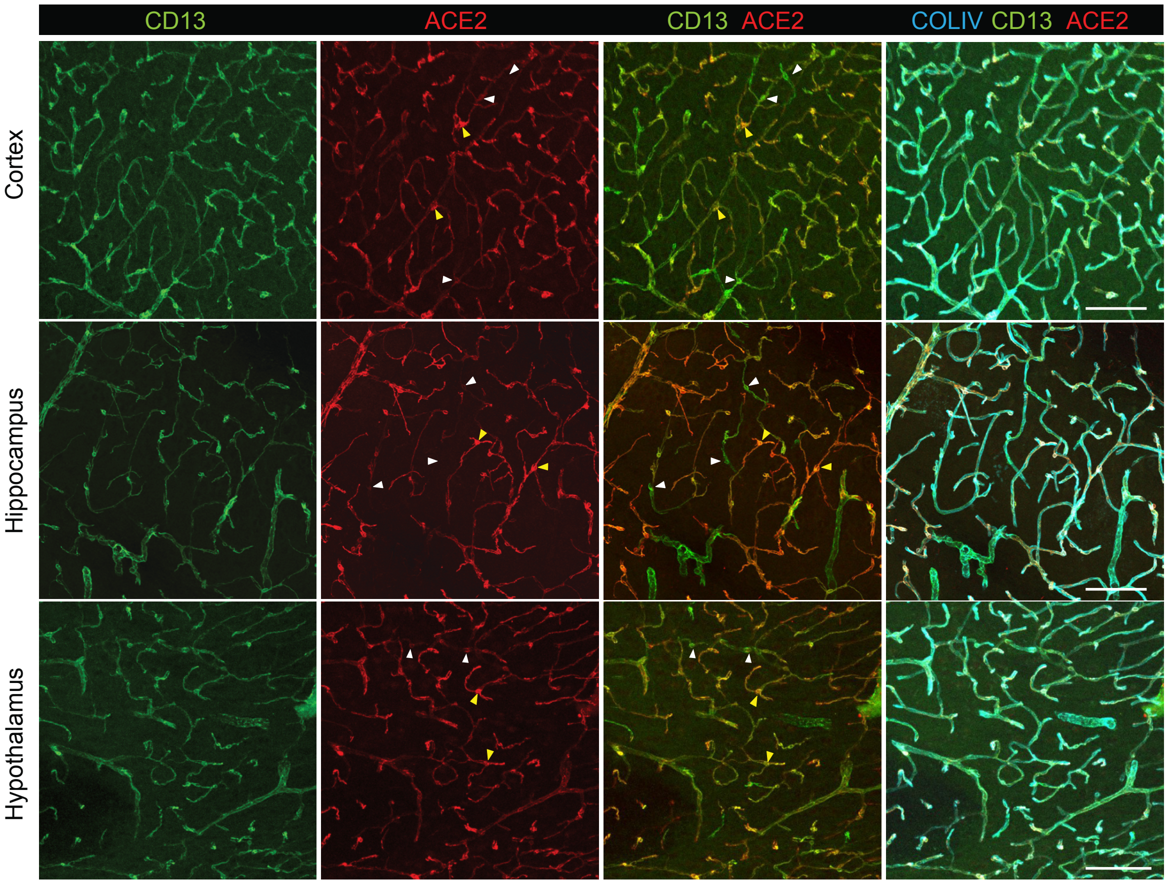
CD13
ACE2
CD13 ACE2
COLIV CD13 ACE2
Cortex
Hippocampus
Hypothalamus

**Figure S8. Age-associated changes in the abundance of ACE2$^{high}$ and ACE2$^{low}$.**
Heterogeneous ACE2 expression (in red) in pericytes (CD13, in green) in the cortex, hippocampus, and hypothalamus of an aged (14–17-month-old) mouse. White arrows indicate ACE2$^{low}$ pericytes and yellow arrowheads indicate ACE2$^{high}$ pericytes. The vascular basement membrane is visualized with collagen IV staining (COLIV, in cyan). Scale bars, 100 µm. Confocal images are presented as maximum intensity projections.

Figure S9 Huang et al.

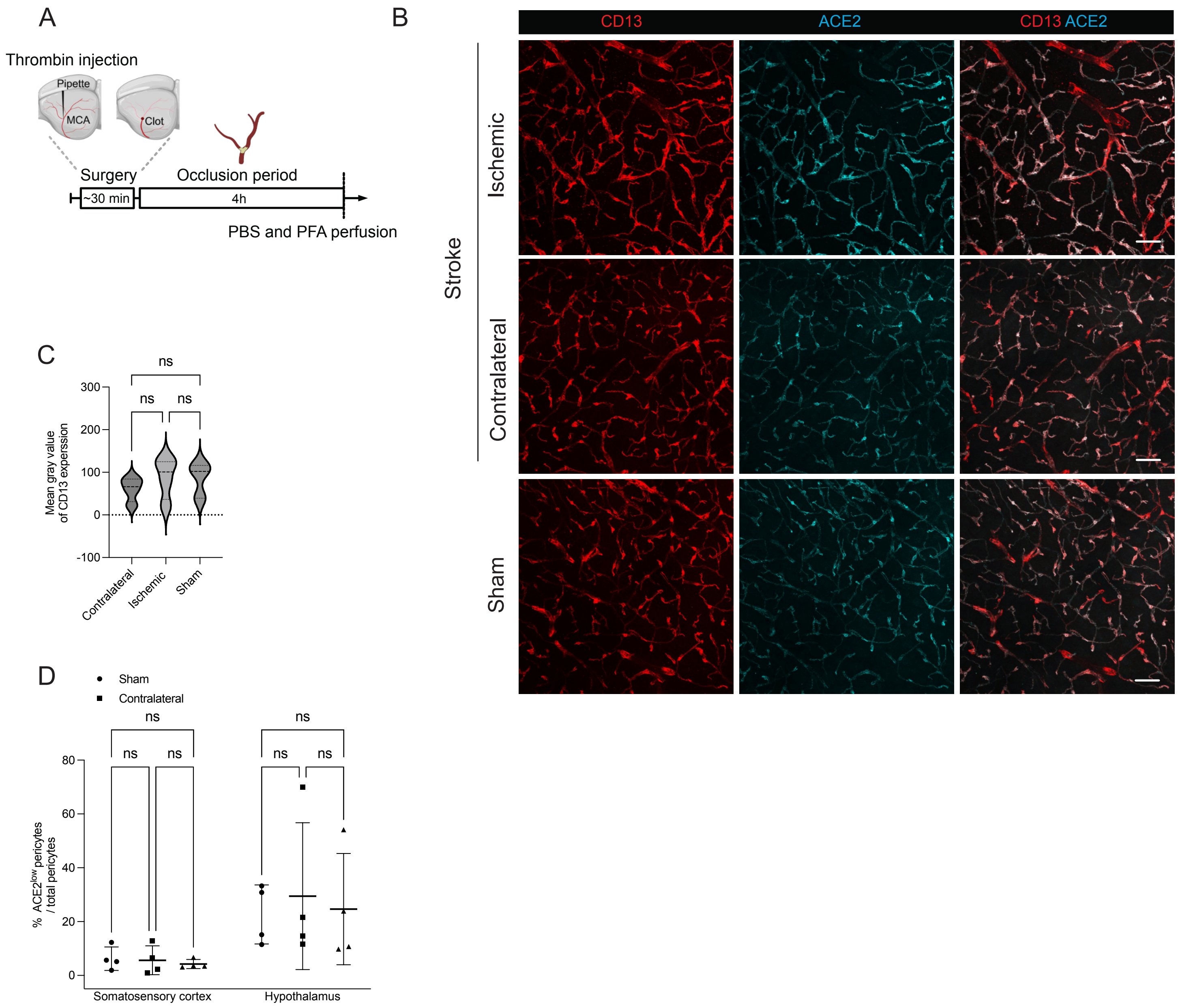

**Figure S9. Acute ischemia does not alter CD13 immunoreactivity or the proportion of ACE2-defined pericyte groups. A** Schematic representation of the transient MCAO experimental workflow. Experimental timeline depicting thrombin-induced middle cerebral artery occlusion (MCAO) in mice. Animals underwent 4 hours of ischemia via thrombin injection. Tissue perfusion and harvesting were performed 4 hours after stroke induction. Surgery took approximately 30 min per animal. **B** Representative images of ACE2 (in cyan) and CD13 (in red) immunostaining in cortical pericytes 4 h after MCAO. Scale bars, 20 µm. **C** Quantification of CD13 immunoreactivity in cortical pericytes 4 h after MCAO. ns, not significant. *n* = 4 mice per group. The thick dotted lines represent the median. The thinner dotted lines show the interquartile range from the 25th to the 75th percentile. One-way ANOVA followed by Tukey's multiple comparison test. **D** Quantification of the proportion of $ACE2^{low}$ pericytes in the somatosensory cortex and hypothalamus of MCAO mice and in the sham-operated mice. Each data point represents an individual animal (*n* = 4 per group). Data presented as mean ± SD. ns, not significant. Two-way ANOVA followed by Tukey's multiple comparison test.

Figure S10 Huang et al.

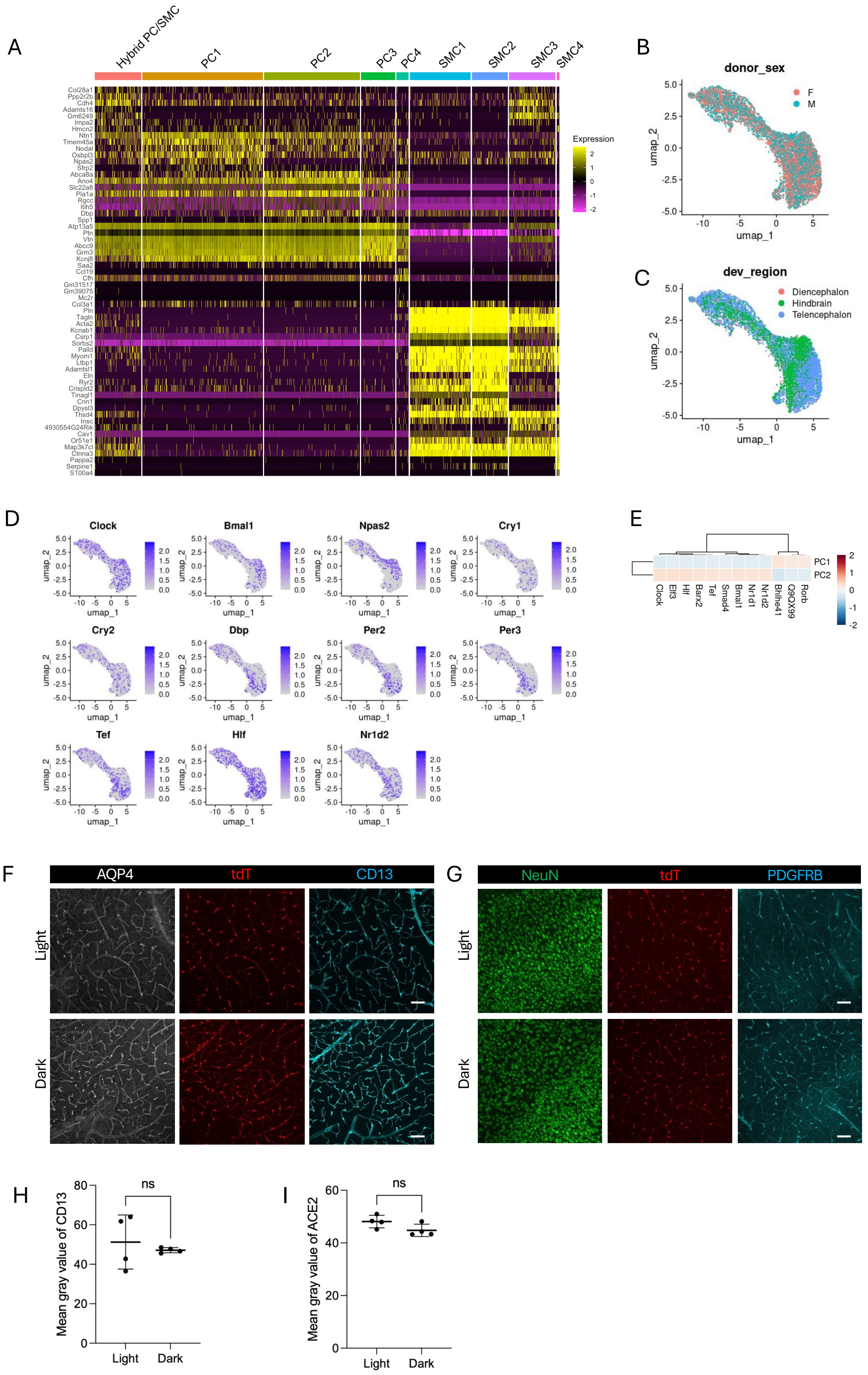

**Figure S10. Transcriptomic analyses and validation of circadian-associated pericyte heterogeneity. A** Heatmap showing the top 10 differentially expressed genes for each cell cluster in the Yao et al. dataset, including cells collected during the light and dark phases. Color intensity reflects scaled average expression (Magenta, low; yellow, high). **B** UMAP visualization of mural cells colored by sex. **C** UMAP visualization of mural cells colored by developmental brain regions as provided in the original metadata. **D** Feature plots showing expression of selected circadian genes within the mural cell cluster. **E** Heatmap showing the top 12 inferred transcription factor activities in PC1 and PC2. Color intensity represents the scaled, inferred TF activity (red: high; blue: low). **F-G** Representative images showing AQP4, CD13, NeuN and PDGFRB expression in brain tissue sections collected during dark and light phases in *Atp13a5*Cre-tdTomato (tdT, in red) mice. Scale bars, 50 µm. **H-I** Quantification of CD13 (I) and ACE2 (J) immunoreactivity in cortical pericytes (tdT, in red) during the light and dark phases in *Atp13a5*Cre-tdTomato mice. *n* =4. Unpaired *t*-test with Welch's correction. ns, not significant. Light phase, ZT4. Dark phase, ZT16.

Figure S11

Huang et al.

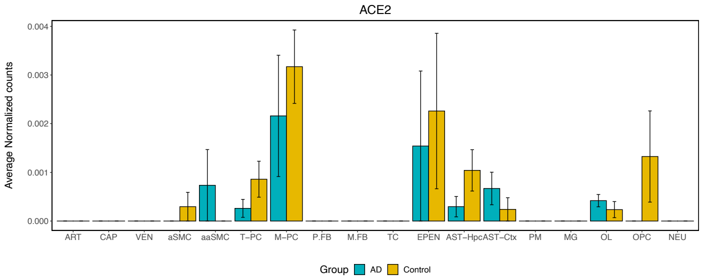

**Figure S11. Heterogeneous *ACE2* expression in human pericytes.** Data are from Yang et al.[14]. ART: Arterial; CAP: Capillary; VEN: Venous; aSMC: Vascular Smooth Muscle Cell; aaSMC: Arteriolar Smooth Muscle Cell; T-PC: Solute transport-Pericyte; M-PC: ECM-regulating Pericyte; P.FB: Perivascular Fibroblast; M.FB: Meningeal Fibroblast; TC: Tcell; EPEN: Ependymal; AST-Hpc: Astrocyte-hippocampus; AST-Ctx: Astrocyte-Cortex; PM: Perivascular Macrophage; MG: Microglia; OL: Oligodendrocyte; OPC: Oligodendrocyte Precursor Cell; NEU: Neuron
